# CCCTC-binding Factor (CTCF) N-terminal Domain Regulates Clustered Protocadherin Gene Expression by Enhancing Cohesin Processivity

**DOI:** 10.1101/2024.08.19.608594

**Authors:** Yijun Zhang, Qiang Wu

## Abstract

CTCF (CCCTC-binding factor) instructs three-dimensional (3D) genome folding by anchoring or forestalling cohesin loop extrusion, but the exact mechanism remains obscure. Here, using clustered protocadherins (*cPcdh*) as model genes, we report that CTCF assists or facilitates cohesin loop extrusion by enhancing its processivity. Specifically, we show that, compared with the *Pcdh α* and *γ* gene clusters, the *Pcdhβ* cluster is greatly affected upon CTCF^Y226A/F228A^ mutation in the N-terminal domain. Given the long-range distance of the *Pcdhβ* cluster from the distal enhancer, this finding has interesting implications in CTCF regulation of cohesin processivity along the linear chromatin during DNA loop extrusion. In particular, the effect on cohesin processivity upon CTCF^Y226A/F228A^ mutation is conspicuously similar to that of WAPL overexpression, suggesting that, in addition to the general view of blocking or forestalling cohesin, CTCF may actually enhance or facilitate cohesin loop extrusion during 3D genome folding. We conclude that CTCF enhances cohesin enrichments via the N-terminal YDF motif in clustered protocadherin genes in a genomic-distance biased manner.

## Introduction

The *cPcdh* locus comprises three closely-linked gene clusters of *α*, *β*, and *γ*, spanning a large region of ∼1 megabase (Mb) genomic DNA in mice and encoding 58 distinct PCDH protein isoforms required for specifying an enormous diversity of neural connectivity in the brain (1,2). The genomic organization of the *Pcdh α* and *γ* clusters is similar, each including a repertoire of large variable exons in a tandem array and a distinct set of three shared downstream constant exons (1,2). Each variable exon is transcribed independently under the control of its own promoter and then spliced to the respective set of downstream constant exons to encode diverse PCDH isoforms. However, members of the *Pcdhβ* cluster each are only made up of a single variable exon without constant ones and thus do not undergo splicing (1,3) (Fig. 1*A*). In addition, all *cPcdh* except c-type genes exhibit stochastic and monoallelic expression patterns in the developing and adult central nervous system (1,4–8). Moreover, the encoded PCDH proteins act as vital cell-surface tags for neural circuit formation, providing individual neurons with a unique molecular ‘identifier’ required for dendritic self-avoidance and axonal tiling in distinct brain regions (7–15). Dysregulation of the *cPcdh* genes is highly relevant to human mental diseases, including autism spectrum disorders (ASD), Down syndrome, and Alzheimer’s disease (AD) (16,17).

**Figure 1.**
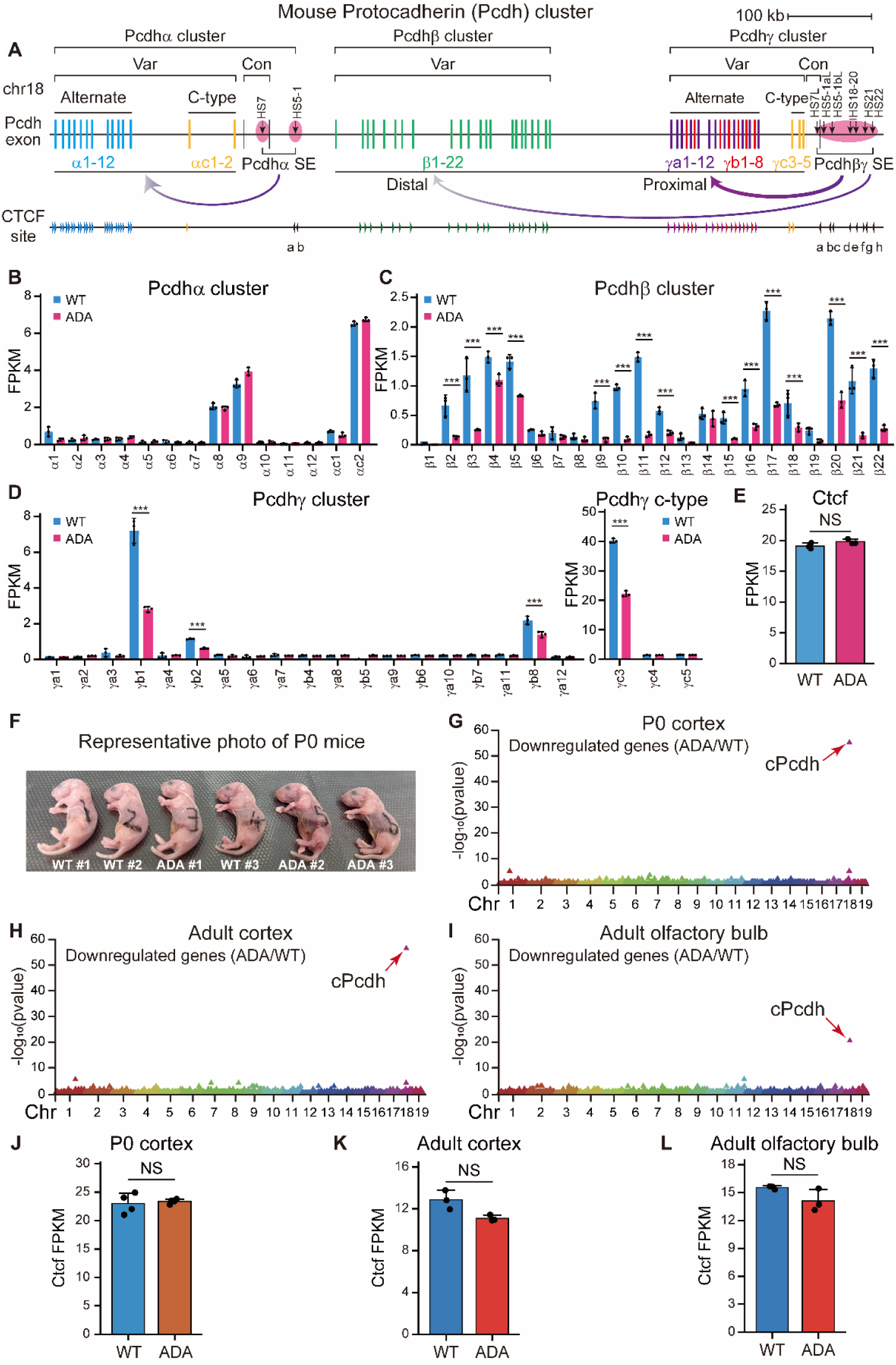
Altered *cPcdh* expression patterns in the CTCF ADA mutant. *A,* Schematic of the mouse *cPcdh* locus. The *Pcdhα* cluster is regulated by a downstream super-enhancer composed of *HS7* and *HS5-1* whereas the *Pcdhβγ* clusters are regulated by a downstream super-enhancer composed of a cluster of enhancers of *HS7L*-*HS22*. The location and orientation of CTCF sites are indicated by arrowheads. Var, variable; Con, constant; SE, super-enhancer. *B*-*D,* RNA-seq of wild type (WT) and CTCF ADA mutant (ADA) Neuro-2a (*N2a*) clones (n=3). *E,* Quantification of the expression levels of the *Ctcf* gene (n=3). *F,* The representative photograph of the WT and heterozygous ADA P0 mice, with their genotypes indicated. *G*-*I,* Manhattan plots showing localized enrichments (using 1Mb^sw^) of downregulated genes in the heterozygous P0 (*G*) and adult (*H*) cortical, as well as olfactory bulb (*I*) tissues, with *cPcdh* as a top-ranking locus. *J*-*L,* Quantitative analyses of the *Ctcf* gene in the heterozygous P0 (n=4) (*J*) and adult (n=3) (*K*) cortical, as well as olfactory bulb (n=3) (*L*) tissues. kb, kilobase. Var, variable. Con, constant. HS, hypersensitive sites. SE, super-enhancer. CTCF, CCCTC-binding factor. FPKM, fragments per kilobase of transcript per million mapped reads. WT, wild-type. ADA, CTCF ADA mutation. NS, non-significant. Chr, chromosome. Data are mean ± S.D from at least three biological replicates, \**P* < 0.05, \*\**P* < 0.01, \*\*\**P* < 0.001; unpaired two-tailed Student’s *t*-test.

The identification of *cis*-regulatory elements (CREs) in the *cPcdh* gene clusters has shed significant insights into the mechanism involved in the *cPcdh* promoter choice. Initial DNase I hypersensitivity assay has characterized several CREs, including a cluster of downstream enhancers of *HS7* and *HS5-1* (super-enhancer) for the *Pcdhα* cluster (18,19). Subsequent studies have identified a large cluster of downstream enhancers of *HS7L*, *HS5-1L*, *HS18-20*, *HS21*, and *HS22* which together constitute a super-enhancer for the *Pcdhβ* and *γ* clusters (6,20–23). Interestingly, members of all three *Pcdh* clusters contain tandem-arrayed CTCF (CCCTC-binding factor) sites in both promoters and enhancers. A striking arrangement of CTCF sites within clustered promoters and enhancers is that the CTCF sites in the promoter regions are in the forward orientation and that those in the enhancer regions are in the reverse orientation, forming convergent CTCF pairs (Fig. 1*A*) (6,21–23). The CTCF/cohesin-mediated long-distance chromatin interactions between distal enhancers and target promoters are the 3D genomic basis of the *cPcdh* promoter choice (6,21–24). Specifically, the tandem directional CTCF sites within the downstream super-enhancers form spatial contacts with the proximal and distal promoter CBS elements through continuous active cohesin “loop extrusion” to erase the genomic-distance biases in favor of stochastic promoter choice (6,8,23). These dynamic cohesin extrusions in the opposite orientation of tandem directional CTCF sites enable them to function as chromatin topological insulators to properly allocate spatial resources for patterned *cPcdh* expression (6). However, the physiological functions of the dynamics of cohesin loop extrusion in the *cPcdh* locus remain poorly understood.

Here, through CRISPR editing of the YDF motif—the key cohesin interacting moiety—of the CTCF N-terminal domain (NTD) in Neuro-2a (*N2a*) cells *in vitro* and in mice *in vivo*, in conjunction with RNA-seq, ChIP-seq, QHR-4C, and Hi-C analyses, we found that members of the *Pcdhβ* cluster are highly susceptible to the perturbed cohesin processivity owing to their distal genomic arrangement relative to the downstream super-enhancer region. We have defined cohesin processivity as the amount of chromatin or size of linear DNA extruded from cohesin, and therefore, we use the genomic distance extruded by cohesin as a surrogate for cohesin processivity. Our data broaden the molecular logic that generates Pcdh isoform diversity required for neural circuit assembly via a unique 3D genome organization.

## Results

### Alteration of *Pcdhβ* expression patterns upon CTCF ADA mutation

Given that CTCF/cohesin-mediated DNA looping between the active promoter and target enhancer is required for the stochastic activation of the *cPcdh* genes (5,21), we investigated how the improper positioning of cohesin affects the promoter choices of the *cPcdh* genes. To this end, we mutated the highly conserved amino acid motif YDF to ADA using DNA-fragment editing through CRISPR single-cell screening in *N2a* cells. Specifically, we screened 166 single-cell clones and obtained two independent homozygous single-cell clones of CTCF ADA mutation (Fig. S1, *A* and *B*).

We performed RNA-seq experiments to assess expression alterations of members of the three *Pcdh* clusters upon CTCF ADA mutation (Fig. 1, *B*-*D*). Remarkably, we observed a significant downregulation in the expression levels of a large repertoire of the *Pcdhβ* cluster (Fig. 1*C*). By contrast, it appears that the effects on the *Pcdhα* and *γ* clusters were less pronounced than those on the *Pcdhβ* cluster (Fig. 1, *B* and *D*). The distinct transcriptional alterations observed within the *cPcdh* locus suggest that members of the *Pcdh β* and *αγ* clusters exhibit varying degrees of sensitivity to CTCF ADA mutation. As a control, we found no significant alteration of *Ctcf* expression levels upon ADA mutation (Fig. 1*E*). Genome-wide expression analyses revealed that ∼5,000 genes exhibited a significant alteration of expression levels upon CTCF ADA mutation. Of note, we observed a higher number of genes downregulated compared to those upregulated (Fig. S1, *C* and *D*).

### Generation of heterozygous *Ctcf^Y226A/F228A^* mice

To further elucidate the crucial role of the YDF motif within the CTCF NTD in regulating cohesin processivity on chromatin *in vivo*, we generated for the first time the *Ctcf^Y226A/F228A^* mice by pronuclear microinjection using the CRISPR/Cas9 system (Fig. S1). Briefly, we designed single guide RNAs (sgRNAs) programming Cas9 to a region adjacent to the genomic sequences encoding the CTCF Y226 and F228 residues. A single-stranded oligodeoxynucleotide (ssODN) donor was co-injected with each sgRNA and Cas9 mRNA to introduce the targeted mutations by homology-directed repair (HDR) (Fig. S1, *E* and *F*). By genotyping 61 P0 chimeric mice, we obtained one mouse with the precise CTCF ADA mutation (Fig. S2*A*) and 14 mice with random indels. The chimeric *Ctcf^Y226A/F228A^* founder was crossed with a wild-type C57BL/6J mouse to generate heterozygous CTCF ADA mutant mice. We obtained two heterozygous *Ctcf^Y226A/F228A^* mice from a total of 5 litters with only 19 pups, suggesting that this CTCF mutation may affect its litter size or fertility (Fig. S2, *B* and *C*). Although the heterozygous *Ctcf^Y226A/F228A^* mice appear to be normal compared to the wild-type littermates (Fig. 1*F* and S2, *D*-*I*), we were unable to obtain homozygous *Ctcf^Y226A/F228A^* mice after repeated breeding attempts. Consequently, we performed all experiments using heterozygous mice.

### Prominent *cPcdh* expression alterations in heterozygous *Ctcf^Y226A/F228A^* mice

We performed RNA-seq experiments using the heterozygous P0 and adult cortical as well as olfactory bulb tissues to further investigate the functional implications of CTCF ADA mutation on gene expression. Consistent with the 3D genome architectural role of CTCF, we found a broad spectrum of genes that display significant expression alterations. Remarkably, 9.3% of the downregulated genes are members of the *Pcdh* clusters, resulting in a unique, ∼231-fold enrichment for downregulated transcripts in the P0 cortex at this locus, as compared to the rest of the genome (*P* = 6.48×10^-56^ by Poisson test; 1-Mb sliding window (1Mb^sw^) applied to n = 302 transcripts) (Fig. 1*G*). There is a similar unique enrichment of *cPcdh* genes for downregulated transcripts in both the adult cortex and olfactory bulb, although the enrichment is less pronounced in the olfactory bulb (Fig. 1, *H* and *I*). As controls, the expression levels of *Ctcf* in the mutant mice are comparable to those in wild-type littermates, suggesting that CTCF ADA mutations do not perturb *Ctcf* expression (Fig. 1, *J*-*L*). Finally, we calculated the proportion of mutated bases of *Ctcf* transcripts and observed that the ratios of mutated bases were comparable to those of wild-type bases (Fig. S1, *G*-*I*).

### CTCF YDF motif regulates quantitative *Pcdhβ* expression

Differential gene expression analyses in the heterozygous P0 cortex, adult cortex, and adult olfactory bulb revealed that among all of the downregulated *cPcdh* genes, the vast majority are members of the *Pcdhβ* cluster (Fig. 2, *A*-*C*). We then calculated expression alterations of each *cPcdh* gene. Remarkably, almost all members of the *Pcdhβ* cluster displayed a significant decrease of expression levels in both the neocortical and olfactory bulb tissues upon CTCF ADA mutation (Fig. 2, *D*-*F*). In addition, members of the *Pcdhβ* cluster exhibited the largest decrease of expression levels compared to the other two clusters in the heterozygous *Ctcf^Y226A/F228A^* mice (Fig. 2, *G*-*I*). This suggests that CTCF ADA mutation has a much stronger effect on members of the *Pcdhβ* cluster than that on those of the *Pcdh α* and *γ* clusters.

**Figure 2.**
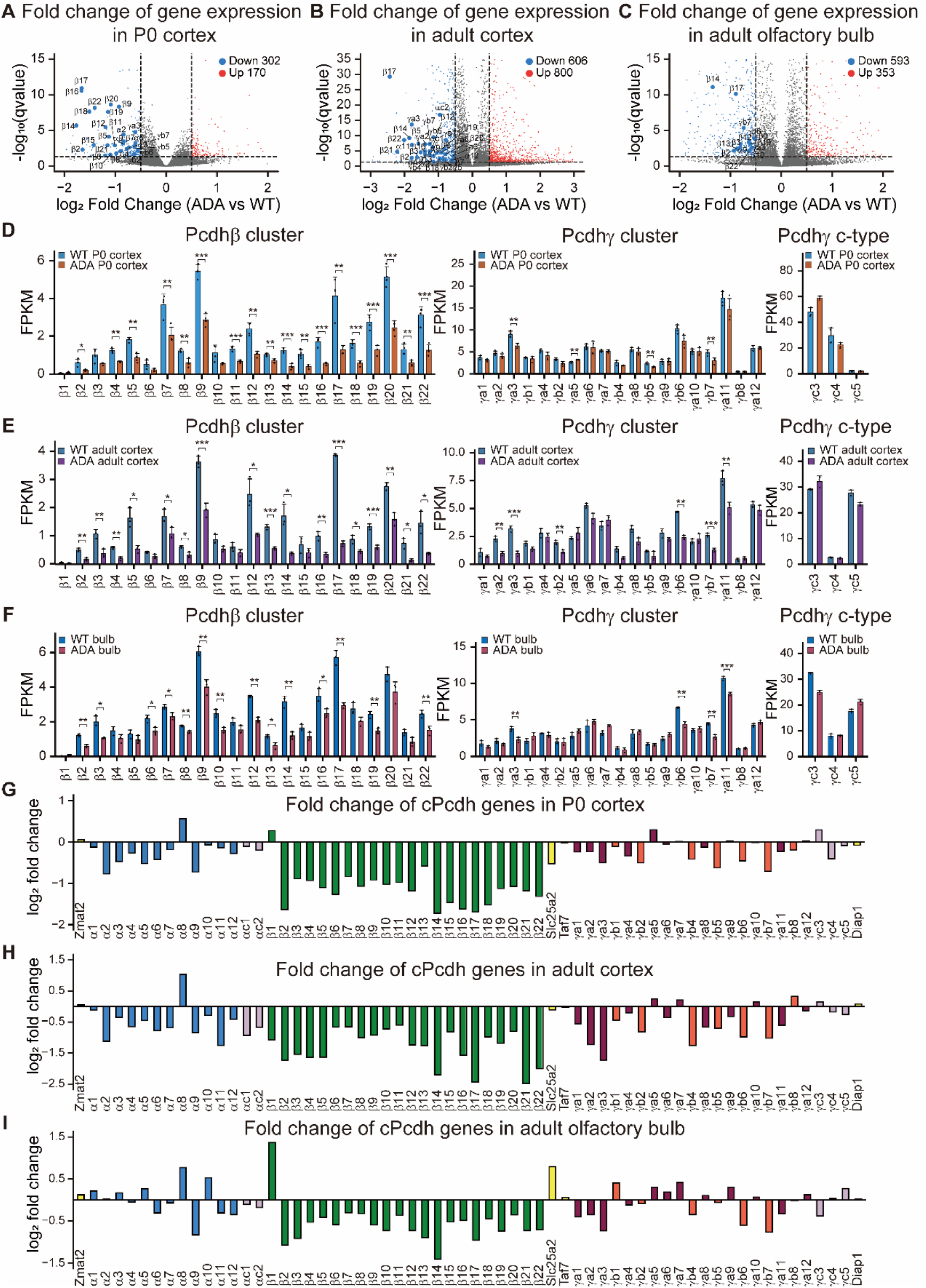
Prominent *Pcdhβ* expression alterations upon CTCF ADA mutation. *A*-*C,* Volcano plots showing all of the downregulated *cPcdh* genes in the heterozygous, compared to their WT littermates, P0 (n=4) (*A*) and adult (n=3) (*B*) cortical, as well as olfactory bulb (n=3) (*C*) tissues. *D*-*F,* Shown are expression alterations of repertoires of the *Pcdh β* and *γ* clusters in the heterozygous P0 (n=4) (*D*) and adult (n=3) (*E*) cortical, as well as olfactory bulb (n=3) (*F*) tissues. *G*-*I,* Bar plots depicting fold changes of gene expression levels of *cPcdh* genes in the heterozygous P0 (n=4) (*G*) and adult (n=3) (*H*) cortical, as well as olfactory bulb (n=3) (*I*) tissues. WT, wild-type. ADA, CTCF ADA mutation. FPKM, fragments per kilobase of transcript per million mapped reads. Data are mean ± S.D from at least three biological replicates, \**P* < 0.05, \*\**P* < 0.01, \*\*\**P* < 0.001; unpaired two-tailed Student’s *t*-test.

### CTCF ADA mutation affects *cPcdh* expression in a genomic distance-biased manner

Increasing evidence has implicated that cohesin-mediated DNA “loop-extrusion” determines the *cPcdh* gene choice via spatiotemporal regulation of long-distance chromatin contacts between distal enhancers and target variable promoters (5,6,8,21,22,25). We asked whether the decreased levels of *cPcdh* gene expression upon CTCF ADA mutation are truly attributed to the chromatin looping mediated by cohesin. To this end, we performed ChIP-seq experiments with a specific antibody against RAD21, a cohesin subunit, in both our cellular and mouse models (Fig. 3 and S3 and S4).

**Figure 3.**
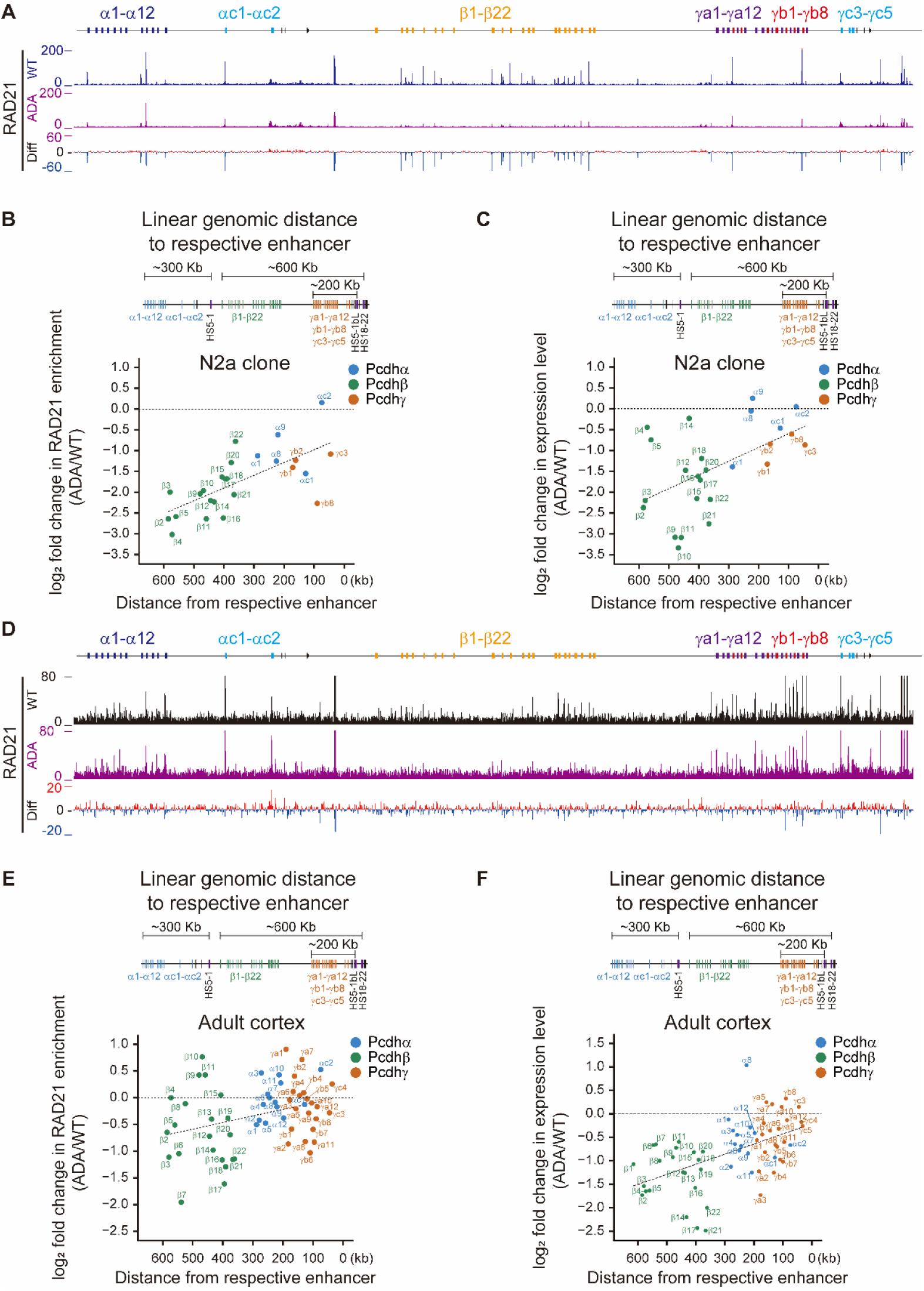
CTCF YDF motif regulates expression levels of *cPcdh* genes in a genomic distance-biased manner. *A,* ChIP-seq profiles of RAD21 at the *cPcdh* locus from WT and CTCF ADA mutant *N2a* clones. *B,* Top: Linear genomic distance of *cPcdh* promoters with respect to their enhancers. Bottom: Scatterplots showing fold changes of RAD21 enrichments at members of three *Pcdh* gene clusters in *N2a* cell clones as a function of the relative linear genomic distance of their promoters to the respective enhancers. *C,* Scatterplots showing fold changes of expression levels of *cPcdh* genes in *N2a* cell clones as a function of relative linear genomic distance between variable promoters and their respective enhancers. *D,* ChIP-seq profiles of RAD21 at the *cPcdh* locus in WT and heterozygous CTCF ADA mice. *E,* Top: Linear genomic distance of *cPcdh* promoters with respect to their enhancers. Bottom: Scatterplots showing fold changes of RAD21 enrichments at the *cPcdh* locus in the heterozygous CTCF ADA mice as a function of the linear genomic distance between target variable promoters and their respective enhancers. *F,* Scatterplots showing fold changes of expression levels of *cPcdh* genes as a function of the linear genomic distance between target variable promoters and their respective enhancers in the heterozygous ADA mice. WT, wild-type. ADA, CTCF ADA mutation. Diff, difference. HS, hypersensitive sites.

Interestingly, we observed a genomic distance-biased decrease of cohesin enrichments for members of *cPcdh* genes, suggesting defects in cohesin processivity upon CTCF ADA mutation (Fig. 3, *A* and *B*). Specifically, in the context of the *Pcdhα* cluster, we found a significant decrease of RAD21 enrichments at *Pcdhα8*, *α9*, and *αc1* (Fig. 3, *A* and *B* and S3), which have CTCF sites associated with their promoters (Fig. 1*A*). This reduction in RAD21 binding at CTCF sites is likely a direct consequence of the CTCF ADA mutation, as the mutated site is a known cohesin interaction site. This decreased cohesin enrichment in the cellular model *in vitro* is similar to previous results in a human HAP1 cell line, suggesting that CTCF stabilizes cohesin-mediated loops through physical interactions (26). By contrast, as an internal control, the RAD21 enrichments remain largely unchanged at the *HS7* enhancer (Fig. S3*E*), which has no associated CTCF site (Fig. 1*A*) (19). In the context of the *Pcdhβγ* clusters, ChIP-seq experiments revealed a significant decrease of the deposition of RAD21 at the super-enhancer region as well as at the variable promoter regions of the *Pcdhβγ* clusters (Fig. 3, *A* and *B* and S3, *F* and *G*). We note that the genomic distance-biased effect is binary for the entire cluster of *β* versus *γ*, but not a graded gene-by-gene decrease within a cluster. In particular, the decrease of RAD21 enrichment at the *Pcdhβ* cluster is much more prominent than that at the *Pcdhα* and *γ* clusters (Fig. S3, *A*-*C*). Interestingly, the genomic distance-biased decrease of cohesin enrichments is consistent with decreased expression levels of members of *cPcdh* genes upon CTCF ADA mutation in *N2a* cell clones (Fig. 3, *B* and *C*).

We observed a similar genomic distance-biased decrease of expression levels of chosen *cPcdh* genes in the heterozygous *Ctcf^Y226A/F228A^* mice *in vivo* (Fig. 3, *D*-*F* and S4). As controls, the enrichment of CTCF itself appears to be unchanged upon CTCF ADA mutation (Fig. S5 and S6). Taken together, these data reveal a genomic distance-biased decrease of *cPcdh* gene expression in both cellular and mouse models.

### CTCF ADA mutation affects cohesin stability on chromatin

To investigate alterations of cohesin-mediated chromatin interactions between distal enhancers and target variable promoters upon CTCF ADA mutation, we performed *in situ* Hi-C experiments and analyzed contact matrices of the *cPcdh* locus. We first evaluated the impact of the ADA mutation on the TAD boundaries using the directionality index algorithm and observed a significant weakening of the strength of *cPcdh* superTAD and subTAD boundaries (Fig. 4*A*) (22,27). In addition, we observed a significant reduction of chromatin contacts within the entire *cPcdh* superTAD upon CTCF ADA mutation (Fig. 4*B*), consistent with the significant decrease of RAD21 enrichments at the *cPcdh* locus (Fig. 3, *D*-*F*). Moreover, there is a significant decrease of long-distance chromatin interactions between the *HS5-1* enhancer and *Pcdhα* variable promoters (Fig. 4*C*). Furthermore, quantitative high-resolution chromosome conformation capture with *HS5-1bL*, *HS18*, or *HS19-20* as an anchor (QHR-4C) revealed a significant decrease of long-range chromatin interactions between the downstream super-enhancer and its target *Pcdhβγ* clusters (Fig. 4*D*). This reduction in long-range enhancer-promoter loops is mainly attributed to the inability of cohesin to extrude long distance of chromatin. Finally, we observed a genomic distance-biased defect in the chromatin contact probability for the *cPcdh* genes upon CTCF ADA mutation (Fig. 4*E*). In particular, although members of both *Pcdh β* and *γ* clusters are activated by the same downstream super-enhancer via cohesin-mediated loop extrusion (6,8,23), the impact on the *Pcdhβ* cluster is much stronger than that on the *Pcdhγ* cluster owing to the greater distance from *Pcdhβ* to the super-enhancer (Fig. 4*E*). Finally, Hi-C analyses revealed that the CTCF ADA mutation mainly affects chromatin loops of large sizes genome-wide (Fig. S7). Together, these data suggest that CTCF may assist distal cohesin enrichment via the NTD YDF motif and that CTCF ADA mutation leads to a cohesin-processivity defect.

**Figure 4.**
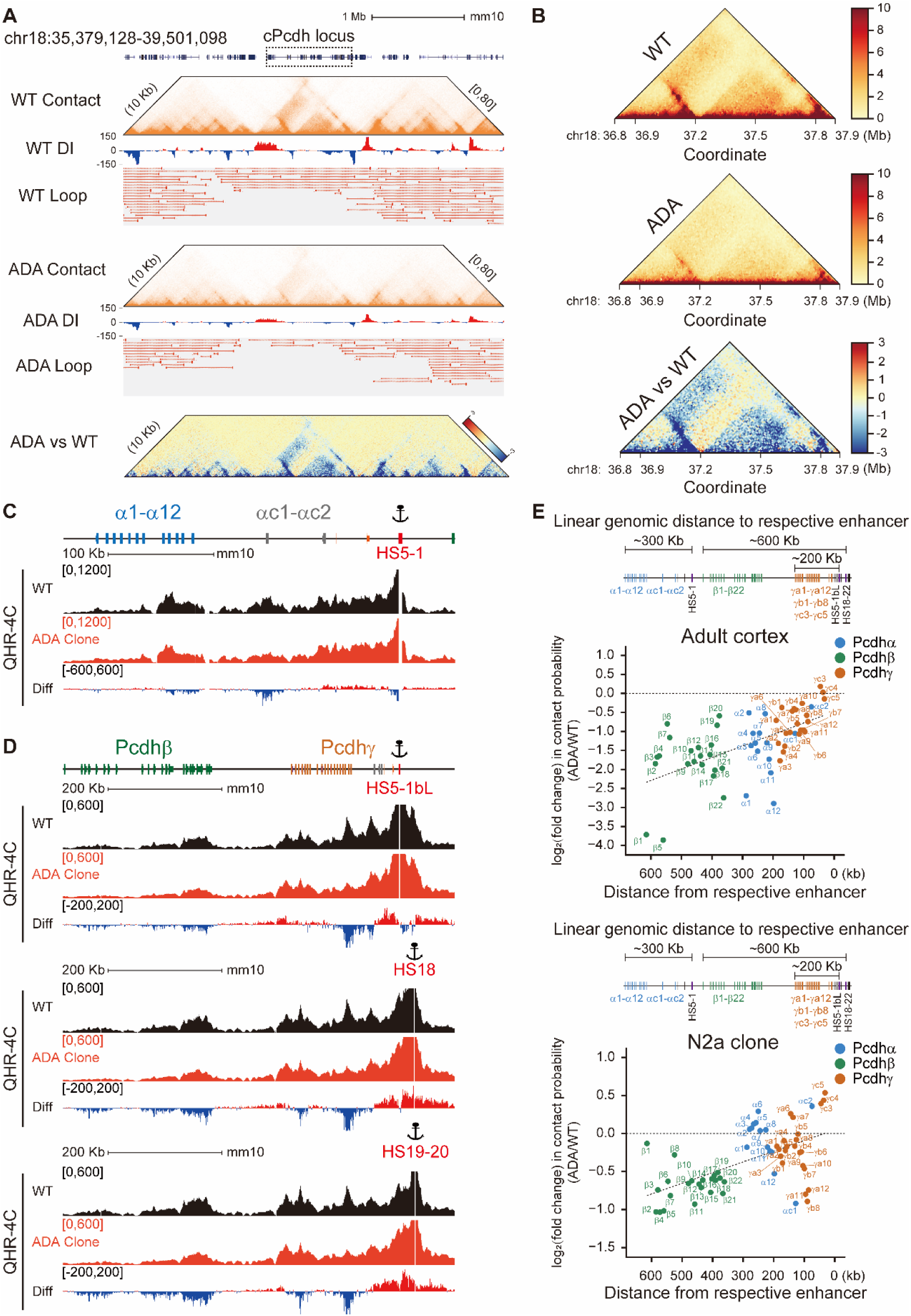
CTCF ADA mutation results in lower cohesin processivity and decreased spatial contact probability between distal promoter-enhancer pairs. *A,* Hi-C contact maps, directionality index scores, and loop intensities around *cPcdh* locus at the chromosome 18 showing a widespread weakening of chromatin loop strength upon CTCF ADA mutation. The differential contact heatmap is also shown. *B,* Close-up of contact maps in the *cPcdh* locus at 10 kb resolution, alongside the differential contact heatmap at the *cPcdh* locus between heterozygous CTCF ADA mutant and WT mice. *C* and *D,* Shown are QHR-4C profiles using *HS5-1*, *HS5-1bL*, *HS18*, or *HS19-20* as an anchor. *E,* Scatterplots depicting fold changes of contact probabilities of *cPcdh* genes as a function of the linear genomic distance between target variable promoters and their respective enhancers in mouse and cellular models. WT, wild-type. ADA, CTCF ADA mutation. DI, directionality index. Mb, megabase. QHR-4C, quantitative high-resolution chromosome conformation capture. Diff, difference. HS, hypersensitive sites. Kb, kilobase.

### Rescue of cohesin releasing by WAPL depletion and CTCF overexpression

To investigate whether CTCF NTD competes with WAPL (wings apart-like protein homolog) for cohesin binding, we generated three WAPL-depleted cell clones in CTCF ADA mutant background (Fig. 5, *A-E* and S8). We next performed QHR-4C experiments with *HS5-1bL*, *HS18*, *HS19-20*, or *HS5-1* as an anchor for each clone and found a significant increase of long-range chromatin interactions between the downstream super-enhancer and its distal target promoters (Fig. 5, *A-D*). In addition, we observed a genomic distance-biased increase in the chromatin contact probability for the *cPcdh* genes upon WAPL depletion (Fig. 5*F*). Consistently, there is a similar genomic distance-biased increase of expression levels of members of *cPcdh* genes (Fig. 5*G*). Finally, we overexpressed wildtype CTCF in the ADA mutant background and also found a similar genomic distance-biased pattern of *cPcdh* gene expression (Fig. 5, *H-J*). This observation suggests that CTCF YDF has an anti-cohesin releasing activity by competing with WAPL.

**Figure 5.**
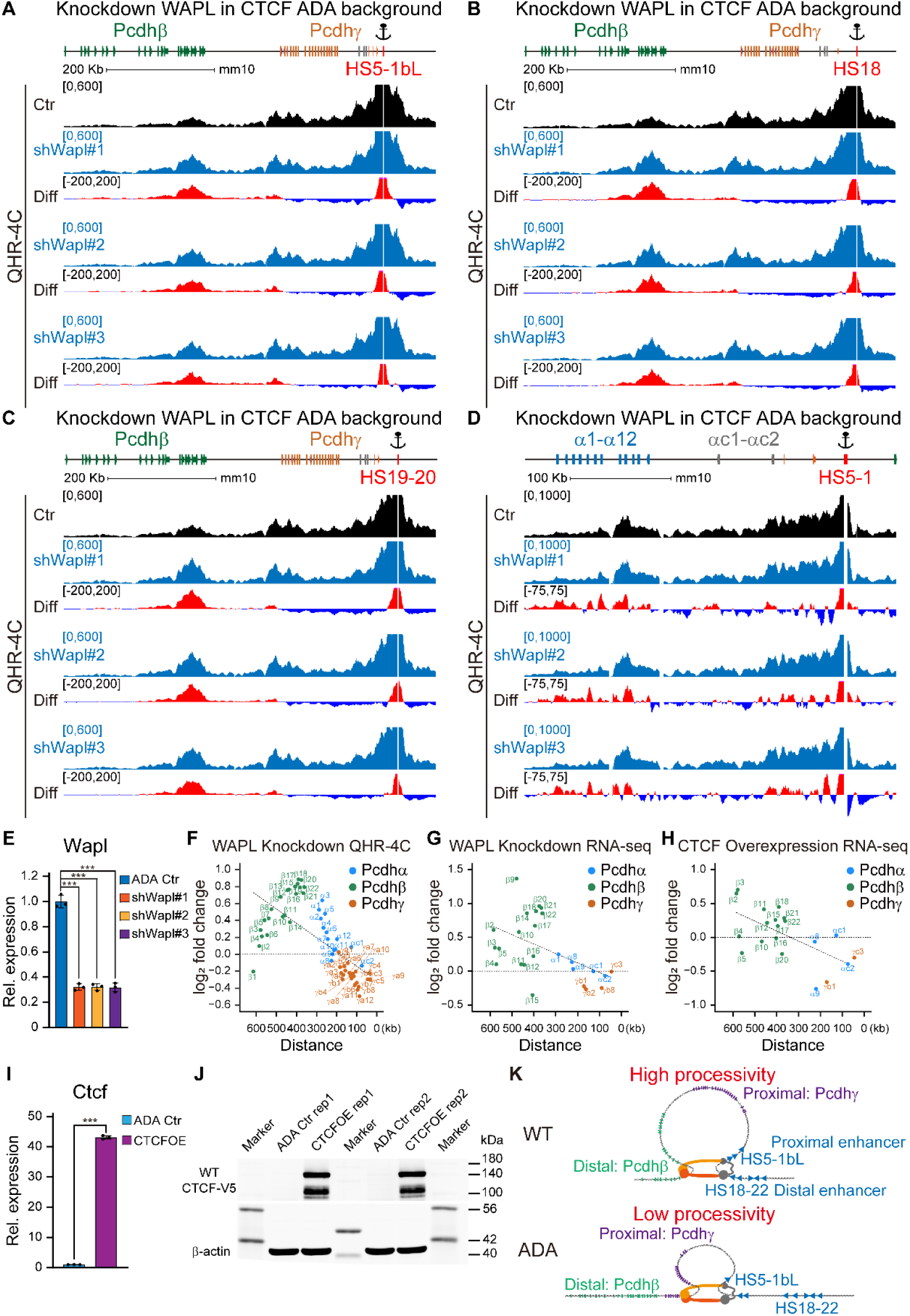
CTCF YDF exerts an anti-cohesin releasing activity by antagonizing WAPL. *A*-*D,* Shown are QHR-4C profiles using *HS5-1bL*, *HS18*, *HS19-20*, or *HS5-1* as an anchor in WAPL-depleted cell clones in the CTCF ADA background. *E,* Quantitative analyses of WAPL expression confirming WAPL knockdown (n=3). *F,* Scatterplots delineating fold changes of contact probabilities of *cPcdh* genes as a function of the linear genomic distance between target variable promoters and their respective enhancers in WAPL-depleted cells. *G* and *H,* Scatterplots illustrating fold changes of expression levels of *cPcdh* genes as a function of relative linear genomic distance between variable promoters and their respective enhancers in WAPL-depleted (*G*) or CTCF-overexpressed (*H*) cells in the CTCF ADA background. *I,* Quantitative analyses of CTCF expression confirming CTCF overexpression (n=3). *J,* Western blot for CTCF overexpression validation. *K,* A model for CTCF regulation of *cPcdh* gene expression patterns in a genomic distance-biased manner. CTCF ADA mutation interferes with its interactions with cohesin, resulting in short residence time on chromatin fibers and low processivity. This leads to prominent dysregulation of members of the *Pcdhβ* cluster. Ctr, control. CTCFOE, CTCF overexpression. kDa, kilodalton. Data are mean ± S.D from three biological replicates, \**P* < 0.05, \*\**P* < 0.01, \*\*\**P* < 0.001; unpaired two-tailed Student’s *t*-test.

## Discussion

### CTCF/cohesin are central for *cPcdh* gene choice

Given that there are approximately 86 billion neurons in the human brain and individual neurons display highly-branched axons and dendrites that intricately connect with one another in diverse brain regions with minimal overlap, it is enigmatic how neurons generate such sophisticated synaptic connectivity to organize precise neural networks for information processing. The cPCDH proteins—a subgroup of cadherin-like cell adhesion molecules—emerge as central candidates for proper wiring of neural circuits (1,7,8,12,13,15,28–31). The CTCF/cohesin-mediated long-distance chromatin contacts between distal super-enhancers and target variable promoters are central for *cPcdh* gene choice during neural development (5,6,8,21–23). Here we have, for the first time, generated a *Ctcf^Y226A/F228A^* mutant mouse model and provided strong genetic evidence that *Ctcf^Y226A/F228A^* mutation severely impacts expression levels of the repertoire of the *cPcdh* genes. In particular, RNA-seq, in conjunction with 4C and Hi-C, showed that members of the *Pcdhβ* cluster are strongly affected in both single-cell *N2a* clones *in vitro* and in heterozygous mice *in vivo* owing to the greatest distance from the super-enhancer. Given the stochastic and monoallelic nature of *cPcdh* gene expression in the brain, it is particularly remarkable that prominent dysregulation of members of the *Pcdhβ* cluster could be observed even in the heterozygous CTCF ADA mutants.

Constitutive CTCF deletion in mice leads to early zygotic lethality, indicating an essential role of CTCF during embryo development (32,33). In addition, conditional CTCF deletion in neurons results in a marked decrease in dendritic arborization and spine density owing to a significant reduction in the expression levels of 53 isoforms of the *cPcdh* genes (34). We observed that the majority of downregulated *cPcdh* genes are members of the *Pcdhβ* cluster upon *Ctcf^Y226A/F228A^* mutation in both mouse and cellular models. This observation is reminiscent of the significant reduction of expression levels of mostly members of the *Pcdhβ* cluster in the heterozygous *Nipbl* deletion mice (35,36). Finally, the regulation of *cPcdh* genes, especially the distal ones, in the brain is related to the SA1-localization at the variable promoters (37).

Emerging CTCF-related neurodevelopmental disorders resulting from numerous *de novo* heterozygous CTCF mutations frequently manifest as syndromic intellectual disability and neurodevelopmental delay such as microcephaly (38–40). In addition, heterozygous mutations in subunits of cohesin complex (SMC1, SMC3, RAD21) or its chromatin loader (NIPBL) cause Cornelia de Lange syndrome (CdLS) which also frequently manifests as intellectual disability and neurodevelopmental delay (41,42). The important role of CTCF/cohesin in regulating *cPcdh* gene expression suggests a neuropathogenic explanation for syndromic intellectual disability from CTCF/cohesin mutations. Indeed, genetic, epigenetic, and 3D genomic mutations and the resulting dysregulation of *cPcdh* gene expression are related to the etiology of numerous neuropsychiatric diseases (16,17). The fact that expression patterns of the *cPcdh* genes are compromised in CTCF ADA mutation, *Nipbl* and *SA1* depletion or mutation, along with the prominent *cPcdh* brain-wiring function, suggest that the dysregulation of *cPcdh* genes resulted from defective cohesin processivity may underlie the pathogenesis of various neurodevelopmental disorders of cohesinopathy.

### CTCF N-terminal domain promotes cohesin processivity

The asymmetrical blocking of extruding cohesin complexes by CTCF at opposite CBS elements in enhancers and promoters leads to the formation of long-distance chromatin loops and determines the promoter choice of *cPcdh* genes (6,21–23,43). Specifically, tandem CTCF sites located at *cPcdh* variable promoters function as chromatin topological insulators to erase the proximity-biased cohesin loop extrusion and enhance the activation of distal promoters (6). In addition, antisense lncRNA transcription leads to DNA demethylation and CTCF deposition and ensures balanced choice of both distal and proximal promoters (5). Moreover, CDCA7 (cell division cycle associated 7) associated DNA methylation loss translates into aberrant upregulation of *cPcdh* genes (44), especially members of the *Pcdhβ* cluster which are distal to the enhancers. The orientation-specific function of CTCF sites are achieved by a combination of its stringent CTCF anti-parallel recognition (45,46) and polar interactions between CTCF NTD and cohesin complex (26,47–49). Given that NTD truncation of CTCF compromises its stalling activity of cohesin loop extrusion and leads to decreased cohesin enrichments at CTCF sites (26,47,48), it is surprising that the ADA mutation does not result in enhanced cohesin processivity. Nevertheless, the compromised cohesin processivity upon CTCF ADA mutation (Fig. 4) is consistent with the role of CTCF in promoting distal promoter-enhancer interactions and the topological insulation function of tandem CTCF sites (6).

Previous studies systematically dissected the enhancer elements of the *Pcdh* gene clusters. The hypersensitive sites *HS5-1* and *HS7*, situated downstream of the *Pcdhα* cluster, were identified as a composite or super-enhancer for the *Pcdhα* cluster (5,8,18,19,21,22,24,25). Additionally, the super-enhancer of *HS5-1bL*, *HS18*, and *HS19-20* predominantly regulates the expression levels of the *Pcdhβ* and *γ* clusters (6,8,20–23). The linear genomic distance of the variable promoters relative to the super-enhancers and CTCF/cohesin-mediated specific promoter-enhancer looping interactions are keys for *cPcdh* gene choice. Interestingly, WAPL dosage in different brain regions functions as a rheostat in regulating *cPcdh* isoform diversity via cohesin processivity (8). Specifically, high WAPL levels in serotonergic neurons lead to low cohesin processivity and promote expression of enhancer-proximal *αc2*; whereas low WAPL levels in the olfactory bulb result in high cohesin processivity, which leads to promoter choice of the enhancer-distal alternate isoforms (8).

We report that the CTCF ADA mutation leads to low cohesin processivity and compromised expression of the members of the *Pcdhβ* cluster, which have the greatest distance from the distal super-enhancer. We propose a model in which the CTCF ADA mutation loses its competition with WAPL for binding to cohesin, resulting in low processivity and compromised expression of the *Pcdhβ* genes (Fig. 5*K*). The YDF motif of CTCF N-terminal domain interacts with the conserved essential surface (CES) of cohesin (26,49). It was initially proposed that the CTCF YDF motif competes with the WAPL N-terminal FGF motifs for binding to cohesin CES (26). However, recent extensive AlphaFold analyses of cohesin subcomplexes suggest that CTCF YDF motif most probably competes with the WAPL C-terminal domain for binding to cohesin CES (50). An alternative mechanism of CTCF in enhancing cohesin processivity could be through interference with the recruitment of WAPL by PDS5 (precocious dissociation of sisters 5), thus compromising the cohesin releasing activity (47,50). Whatever the underlying mechanism, our data suggest CTCF YDF has an anti-cohesin releasing activity by competing with WAPL. The exact mechanism by which CTCF and WAPL regulate cohesin processivity or chromatin association awaits future studies.

### Experimental procedures

#### Cell culture

Mouse Neuro-2a (*N2a*) cells were cultured in a modified Eagle’s medium (Gibco 11095080) supplemented with 10% fetal bovine serum (FBS) (Sigma-Aldrich F0193), 1% penicillin-streptomycin (Gibco 15140122), and 1% Non-Essential Amino Acids (Gibco 11140050). Typically, *N2a* cells were incubated at 37°C in a humidified incubator with 5% (v/v) CO_2_.

Human embryonic kidney cells (*HEK293T*) were cultured in Dulbecco’s modified Eagle’s medium (Gibco 11995065) supplemented with 10% fetal bovine serum (FBS) (Sigma-Aldrich F0193) and 1% penicillin-streptomycin (Gibco 15140122). The cells were typically maintained at 37°C in a humidified incubator with 5% (v/v) CO_2_ for generating high titers of lentiviral particle for WAPL depletion.

For cell passaging, when cells reached about 80% confluency, the culture medium was aspirated from the cell culture dish. The cells were then washed gently once with 1 ml of 1 × PBS (Gibco 70011044) (*Table S2*) to remove residual medium and cell debris. 1 ml of 0.25% trypsin-EDTA (Gibco 25200056) was added, followed by incubation at 37°C for 3 min to detach cells from the dish surface. Next, 1 ml of cell culture medium was added to neutralize the trypsin. The cell suspension is then transferred to a new 15-ml centrifuge tube and centrifuged at 900 rpm for 5 min at room temperature. The supernatant was discarded, and 1 ml of fresh medium was added to resuspend the cell pellet. Finally, cells are sub-cultured via transferring 250 μl of cell suspension into a new dish containing 5 ml of fresh culture medium.

#### Plasmid construction

The construction of plasmids (*Table S1*) for the generation of CTCF ADA mutant (ADA) Neuro-2a (*N2a*) clones was carried out as previously described (51,52). In brief, to generate constructs for sgRNA transcription, 2 μl each of two complementary oligonucleotides (Mut_sgRNA_F and Mut_sgRNA_R) (*Table S1*) at a concentration of 20 μM and 2 μl of NEBuffer 2 (NEB B7002S) were added into an Eppendorf tube, and the reaction volume was adjusted to 20 μl with nuclease-free water. The reaction mixture was heated in boiling water for 1 min, then allowed to gradually cool to room temperature to form double-stranded DNA (dsDNA) with 5′ overhangs of ‘ACCG’ and ‘AAAC’. In addition, 1 μl of *Bsa*I (NEB R3733S) was added to another Eppendorf tube containing 3 μg of *pGL3* plasmid at a total volume of 50 μl, and the mixture was incubated at 37°C for 2 h to obtain a *Bsa*I-linearized *pGL3* vector. Finally, the annealed dsDNA is then cloned into a *Bsa*I-linearized *pGL3* vector with *U6* promoter to enable sgRNA transcription with the U6 polymerase.

For construction of donor plasmids used in homologous recombination, the left homologous arm of 883-bp was generated by PCR (95°C, 3 min; 95°C, 15 s, 58°C, 15 s, 72°C, 1 min for 35 cycles; and a final extension at 72°C, 5 min) from total mouse genomic DNA with a forward primer (Left_Mut_F) (*Table S1*) containing vector backbone sequences and a reverse primer (Left_Mut_R) (*Table S1*) containing the designed CTCF ADA mutation and a synonymous mutation at its 5’ end. In addition, the right homologous arm of 885-bp were generated by PCR (95°C, 3 min; 95°C, 15 s, 60°C, 15 s, 72°C, 1 min for 35 cycles; and a final extension at 72°C, 5 min) from total mouse genomic DNA with a forward primer (Right_Mut_F) (*Table S1*) containing the same mutations and a reverse primer (Right_Mut_R) (*Table S1*) also containing vector backbone sequences. Moreover, to obtain the vector backbone, 1 μl of *Eco*RI (NEB R3101S) and 1 μl of *Hin*dIII (NEB R3104S) were added to an Eppendorf tube containing 2 μg of *pUC19* plasmid at a total volume of 50 μl, and the mixture was incubated at 37°C for 2 h to obtain a linearized *pUC19* vector. To construct the donor plasmid, the two homologous arms with designed mutations and the linearized vector backbone were joined using the Gibson Assembly (Vazyme C113) according to the protocol recommended by the manufacturer. The oligonucleotides for plasmid construction were listed in *Table S1*. All plasmids were validated by Sanger sequencing.

#### CRISPR screening of CTCF ADA single-cell clones

The generation of CTCF ADA single-cell clones was performed using CRISPR/Cas9-mediated DNA fragment editing (51,52). Initially, cells are seeded in a 6-cm culture dish and allowed to reach ∼80% confluency. The mixture containing 2 μg of the Cas9 plasmid, 1 μg of the sgRNA plasmid, and 1 μg of the donor plasmid, along with 10 μl of transfection reagent P3000 (Invitrogen L3000015) and 100 μl of MEM (Gibco 11095080) was gently mixed with another mixture consisting of 100 μl of MEM (Gibco 11095080) and 10 μl of Lipo3000 (Invitrogen L3000015). Mixtures were then incubated at room temperature for 15 min. Subsequently, the transfection mixture was added to the cells, and the cells were continuously incubated for additional 2 days. The medium was replaced with fresh MEM each day to remove harmful substances and cell debris. Two days after transfection, puromycin (Solarbio P8230-25mg) was added to the cell growth medium at a concentration of 2 μg/ml to initiate the antibiotic selection for 4 days. Finally, the cells are cultured for an additional week in antibiotic-free medium to allow for recovery, before being used for screening single-cell clones.

To screen single-cell clones, the cultured cell mixture was transferred to a new tube and centrifuged at 2,000 × g to pellet the cells. The supernatant was discarded, and the cell pellets were lysed in 30 μl of Solution A (25 mM NaOH) at 95°C for 20 min and neutralized with 30 μl of Solution B (25 mM Tris-HCl pH 6.8). The genomic DNA of the cultured cell mixture was then screened for the targeted mutations by PCR (95°C, 3 min; 95°C, 15 s, 58°C, 15 s, 72°C, 25 s for 35 cycles; and a final extension at 72°C, 5 min) using specific primers with the 3’ end of the forward primer matching the designed mutation and a synonymous mutation (geno_F1_Mut and geno_R1) (*Table S1*). The cell mixture with the targeted mutations was diluted and seeded into eight 96-well culture plates, with approximately one cell per well. The plates were then cultured for a period of 14 days to let the single cell to grow into a colony. The colonies were picked under a microscope and genotyped by PCR (95°C, 3 min; 95°C, 15 s, 58°C, 15 s, 72°C, 25 s for 35 cycles; and a final extension at 72°C, 5 min) using specific primers (primer pairs: geno_F1_Wt and geno_R1 for WT band detection; primer pairs: geno_F1_Mut and geno_R1 for CTCF ADA band detection) (*Table S1*).

We screened a total of 166 clones and obtained two homozygous CTCF ADA *N2a* clones. The homozygous CTCF ADA single-cell clones were validated by Sanger sequencing. The primer sequences used for CRISPR DNA-fragment editing were listed in *Table S1*.

#### Lentivirus packaging for the establishment of stable cell lines with *Wapl* depletion

The plasmid construction (*Table S1*) for efficient *Wapl* knockdown was performed following previous protocols (51,52). Briefly, to generate constructs for shRNA transcription, complementary oligonucleotides (shCtr_F and shCtr_R) (*Table S1*) were annealed to form double-stranded DNA (dsDNA) with 5′ overhangs of ‘CCGG’ and ‘AATT’. In addition, the *pLKO.1* plasmid was linearized using *Age*I (NEB R3552S) and *Eco*RI (NEB R3101S). Subsequently, the annealed DNA was cloned into the linearized *pLKO.1* vector under the U6 promoter for shRNA transcription.

To establish stable cell lines with efficient *Wapl* knockdown, a mixture of the *pLKO.1* lentivirus transfer plasmid, packaging helper plasmid (psPAX2, Addgene 12260), and envelope helper plasmid (pMD2.G, Addgene 12259) was co-transfected into *HEK293T* cells using Lipofectamine 3000 reagents (Invitrogen L3000015) to produce high-titer lentivirus particles. After 24 hours of incubation, the medium was replaced with fresh complete medium containing 30% FBS. Lentivirus particles were harvested from the HEK293T supernatant after 48 and 72 hours of transient transfection, and subsequently used for lentivirus transduction in *N2a* cells to generate stable cell lines.

#### Transient transfection in *N2a* cells to achieve CTCF overexpression

Extraction of total RNA was conducted with Trizol reagents (Invitrogen 15596018), followed by reverse transcription using 1 μg of total RNA. In brief, the residual genomic DNA was digested using 4 μl of gDNA wiper mix (Vazyme R312), and the first-strand cDNA synthesis mixture was incubated at 37°C for 45 min to initiate reverse transcription.

To clone the CTCF coding sequence for overexpression, the mouse cDNA library was diluted and used as the template to amplify the CTCF coding sequence using specific primers (mCtcf_EcoRI_F and mCtcf_NotI_R). Additionally, to obtain the vector backbone, 1 μl of *Eco*RI (NEB R3101S) and 1 μl of *Not*I (NEB R3189S) were added to an Eppendorf tube containing 2 μg of *pLVX* plasmid at a total volume of 50 μl. The mixture was then incubated at 37°C for 4 h to obtain a linearized *pLVX* vector. To construct the overexpression plasmid, the CDS construct and the linearized backbone were joined using T4 DNA ligase (NEB M0202S).

CTCF ADA mutant (ADA) *N2a* cell clones growing at 60% confluency in 6-well plates were transiently transfected with 2.5 μg of *pLVX* plasmids using Lipofectamine 3000 reagents (Invitrogen L3000015) following the manufacturer’s protocol. After 48 h, the transfected *N2a* cells were subjected to antibiotic selection with 2 μg/ml puromycin in culture medium for 48 h. Survived cells were then cultured for an additional 24 h in puromycin-free culture medium before being harvested for RNA-seq and Western blot experiments.

#### Western blot

For protein extraction, transiently transfected cells were lysed using RIPA lysis buffer (50 mM Tris-HCl pH 7.4, 150 mM NaCl, 1% Triton X-100, 1% sodium deoxycholate, 0.1% SDS, and 1 × protease inhibitors). The total protein was denatured at 95°C for 10 min, separated by SDS-PAGE, and transferred to nitrocellulose membranes. The membrane was blocked using 5% nonfat milk in 1 × TBST and then incubated overnight at 4°C with a primary antibody. After three washes with 1 × TBST, the membrane was incubated at room temperature for 1 h with a secondary antibody, followed by scanning using the Odyssey System (LI-COR Biosciences).

#### Synthesis of the sgRNA and Cas9 mRNA for microinjection by *in vitro* transcription

The generation of CRISPR components was performed as previously described (6,23). Briefly, the sgRNA targeting sequence was systematically selected using a computational program (53). The template for sgRNA *in vitro* transcription was generated by PCR with a pair of primers. The forward primer (T7_Mut-sgRNA-F1 or T7_Mut-sgRNA-F2) (*Table S1*) includes a T7 promoter sequence (5’-TAATACGACTCACTATA-3’), the sgRNA-target sequence, and the sgRNA scaffold sequence (5’-GTTTTAGAGCTAGAAATAG-3’) (Sangon). The common reverse primer (sgRNA-R) matches the sgRNA backbone (*Table S1*). The total volume of PCR reaction is 50 μl, containing 1 μl of forward primer, 1 μl of common reverse primer, 25 μl of 2 × Phanta Buffer (Vazyme P505-d1), 1 μl of dNTP (Vazyme P505-d1), 1 μl of Phanta (Vazyme P505-d1), and 10 ng of the *pGL3* plasmid as a template, to amplify (95°C, 3 min; 95°C, 20 s, 63°C, 25 s, 72°C, 10 s for 35 cycles; and a final extension at 72°C, 1 min) a linear DNA template for sgRNA *in vitro* transcription.

The PCR product was purified using the PCR purification kit (Qiagen 28104) and eluted in 30 μl of nuclease-free water. The volume of the purified product was brought to 400 μl with RNase-free water (Invitrogen 10977023), followed by the addition of 400 μl of phenol-chloroform (ACMEC AC13309) to remove RNase. The sample was then vigorously vortexed to mix thoroughly and centrifuged at 17,000 × g for 15 min at room temperature. DNA was precipitated by transferring 320 μl of supernatant to a new RNase-free microcentrifuge tube and adding 32 μl of 3 M sodium acetate (pH 5.5) (Invitrogen AM9740), 2.5 μl of RNA-grade glycogen (Thermo Scientific R0551), 640 μl of ice-cold ethanol. The sample was vortexed and then stored at −80°C for at least 30 min. The precipitated DNA was pelleted by centrifuging at 12,000 × g for 20 min at 4°C and washed with 1 ml of freshly-made 75% ice-cold ethanol. Finally, the washed sample was then centrifuged at 12,000 × g for 5 min at 4°C, and the DNA pellet was resuspended in 30 μl of RNase-free water (Invitrogen 10977023).

The sgRNA *in vitro* transcription was performed using the MEGAshortscript Kit (Invitrogen AM1354) following the protocol recommended by the manufacturer. In brief, 500 ng of purified product was mixed with 2 μl each of ATP, CTP, GTP, and UTP, along with 2 μl of 10 × Buffer and 2 μl of T7 RNA polymerase (Invitrogen AM1354). The sample was mixed gently and incubated at 37°C for 3 h, followed by the addition of 1 μl of TURBO DNase (Invitrogen AM1354). It was then incubated at 37°C for an additional hour. The transcribed product was brought to a total volume of 100 μl with RNase-free water (Invitrogen 10977023) to purify the RNA. The sgRNA was recovered using the MEGAclear Kit (Invitrogen AM1908) according to the protocol recommended by the manufacturer. In brief, the sample was gently mixed with 350 μl of Binding buffer (Invitrogen AM1908), followed by the addition of 250 μl of ethanol and mixing. The sample was then applied to the filter column (Invitrogen AM1908) and left to stand by 5 min, followed by centrifugation at 14,000 × g for 1 min. After discarding the flow-through, the column was washed twice with 500 μl of Washing solution (Invitrogen AM1908) at 14,000 × g for 1 min. After discarding the Washing solution, the column was centrifuged at 14,000 × g for 1 min to completely remove the Washing solution. Finally, the sgRNA was eluted from the filter with 30μl of Elution solution (containing 0.2 mM EDTA) at 14,000 × g for 1 min.

The concentration of the purified RNA was measured using NanoDrop 2000 Spectrophotometer. The A260/A280 ratio of high-quality RNA is approximately 2.0, while the A260/A230 ratio is at least 2.2. The sgRNA was then divided into aliquots of 2.5 μg per tube and stored at −80°C to avoid freeze-thaw cycles. The primer sequences used for the sgRNA generation were listed in *Table S1*.

To synthesis the Cas9 mRNA, 3 μl of *Xba*I (NEB R0145S) was used to digest 6 μg of the *pcDNA3.1-Cas9* vector, which can express Cas9 under control of the *T7* promoter, in a total volume of 100 μl to obtain a linearized vector. The linearized vector was purified through a gel extraction column (Qiagen 28704). The linearized product was eluted in 30 μl of nuclease-free water and brought to 400 μl with RNase-free water, followed by the addition of 400 μl of phenol-chloroform (ACMEC AC13309) to completely remove RNase. The sample was vigorously shake until the solution turns white and centrifuged at 17,000 × g for 15 min at room temperature. After ethanol precipitation, the DNA pellet was resuspended in 15 μl of RNase-free water.

The Cas9 mRNA was transcribed *in vitro* from the purified linearized product using the mMACHINE T7 ULTRA kit (Invitrogen AM1345) according to the manufacturer’s instructions. Briefly, 1 μg of *Xba*I-linearized product was mixed with 10 μl of 2 × NTP/ARCA, 2 μl of 10 × Buffer, and 2 μl of T7 RNA polymerase (Invitrogen AM1345) in a total volume of 20 μl. The sample was then incubated at 37°C for 1 h, followed by the addition of 1 μl of TURBO DNase (Invitrogen AM1345) and incubation at 37°C for 1 h. The transcription reaction was added with a poly(A) tailing mixture containing 36 μl of RNase-free water, 20 μl of 5 × *E*-PAP Buffer, 10 μl of 25 mM MnCl_2_, and 10 μl of ATP solution, and 4 μl of *E*-PAP enzyme (Invitrogen AM1345) at a total volume of 100 μl. The tailing reaction was incubated at 37°C for 45 min. The Cas9 mRNA was also recovered using the MEGAclear Kit (Invitrogen AM1908). The concentration of the purified RNA was measured using the spectrophotometer (NanoDrop 2000). Finally, the Cas9 mRNA was divided into aliquots of 5 μg per tube and stored at −80°C to avoid freeze-thaw cycles.

#### Generation of CTCF ADA mutation mice by homologous recombination using CRISPR/Cas9 system via microinjection

C57/BL6J and Institute of Cancer Research (ICR) mice were maintained at 23°C in an SPF (specific pathogen free) facility following a 12-h (7:00‒19:00) light and 12-h (19:00‒7:00) dark schedule. All animal experimental procedures were approved by the Institutional Animal Care and Use Committee (IACUC) of Shanghai Jiao Tong University (protocol#: A2016041). Briefly, for mice superovulation, the Pregnant mare serum gonadotropin (PMSG) (Solarbio P9970) or human chorionic gonadotropin (hCG) (MCE HY-107953), supplied as a lyophilized powder, is resuspended at a final concentration of 100 U/ml using 0.9% sterile saline (Sigma-Aldrich S3014) solution and then divided into aliquots of 1 ml per tube. The optimal age of mice prepared for superovulation lies between 3 and 6 weeks. Twelve mice were induced to super-ovulate by intraperitoneal injection of 10 U of PMSG on day 1 at 3 pm. On day 3 at 1 pm, 10 U of hCG was administrated. After the injection of hCG, each female mouse is placed in a cage with a stud male for crossing.

Next morning, the females were checked for a copulation plug, and usually 7-10 out of 12 females will be plugged. The unplugged superovulated females can be used for a second time. Typically, about 25 zygotes can be obtained from a 3- to 6-week-old superovulated C57/BL6J, yielding a total of ∼200 zygotes for microinjection. The plugged mice were culled by cervical dislocation, followed by dissection of the oviduct and transference of the oviduct into pre-warm M2 medium containing hyaluronidase (Sigma-Aldrich H4272-30MG) at a final concentration of 0.3 mg/ml. The zygotes were pipetted up and down a few times using a mouth-controlled glass capillary until the cumulus cells fell off. The zygotes were then transferred to a droplet containing no hyaluronidase (Sigma-Aldrich H4272-30MG) to rinse off the hyaluronidase solution and cell debris. The zygotes can be kept at 37°C in a humidified incubator with 5% (v/v) CO_2_ until needed. 15-20 zygotes were placed into an M2 medium droplet in a chamber for microinjection.

To prepare a chamber for microinjection, the M2 medium droplets were set up the day before the zygote collection to allow for gas equilibrium at 37°C in a humidified incubator with 5% (v/v) CO_2_. Briefly, 40-μl drops of M2 medium were dispensed on the bottom of a 35-mm culture dish. The droplets were flooded with 3 ml of mineral oil (Sigma-Aldrich M5310). In addition, another 35-mm culture dish containing 3 drops of 25-μl each was used as an injection chamber, which was flooded with 2.5 ml of mineral oil (Sigma-Aldrich M5310) and equilibrized at 37°C in a humidified incubator with 5% (v/v) CO_2_.

The injection mixture containing 5 μg of Cas9 mRNA, 2.5 μg of sgRNA, and 10 μg of ssODN was mixed in advance at a total volume of 50 μl and was divided into aliquots of 10 μl per tube. The mixture was loaded into the injection capillary using a mouth-controlled glass capillary and injected into the zygote at a very low pressure, delivering 2 picoliters. Once all of the zygotes in the injection chamber were injected, they should be immediately transferred back into a droplet of M2 medium (Sigma-Aldrich MR-015), which is flooded with mineral oil (Sigma-Aldrich M5310), and incubated at 37°C/5% CO_2_ incubator for 30 min to recover. Only zygotes exhibiting normal morphology were implanted into the infundibulum of surrogate pseudo-pregnant ICR females.

For zygotes implantation, the surrogate pseudo-pregnant ICR females were weighed and anesthetized by intraperitoneal injection, and placed on the lid of a 10-cm dish. The back of the mouse was wiped with tissues soaked in 75% ethanol. The fur around the dorsal midline was removed using fine scissors. A single incision was made along the dorsal midline, perpendicular to the vertebral column, just below where the last rib ended. A smaller incision in the body wall was made to expose the reproductive tract. The fat pads were pulled out, and the ovary, oviduct, and uterus came with the fat pads. The microinjected zygotes were then loaded into the transfer pipette. First, a small amount of M2 medium was taken up into the pipette, followed by an air bubble, then M2 medium, and then a second air bubble. The zygotes were drawn up in a minimal volume of M2 medium, followed by an additional small bubble at the tip of the pipette. The infundibulum of the oviduct was gently cradled by blunt forceps, and the prepared transfer pipette was inserted into the infundibulum. The zygotes were blown into the ampulla, and the reproductive tract was put back into the body cavity. Finally, the implanted ICR females were then placed into a clean cage on a warming plate until they recovered from the anesthetic.

To obtain surrogate pseudo-pregnant ICR mice for zygote implantation, the females in estrus were mated with the sterile males the day before implantation. To prepare sterile ICR males, 8-week-old ICR males were chosen to undergo the vasectomy operation. The males were weighed and anesthetized by intraperitoneal injection. A small 15-mm incision was made along the midline of the abdomen using fine scissors. A similar-sized incision in the body wall was made to expose the tissue. The testicular fat pads were pulled out, and the testis, vas deferens, and epididymis came with the fat pads. The vas deferens was held by fine forceps and cauterized with the red-hot tips of another forceps. Next, the testis was carefully placed back inside the body cavity. The same procedures were repeated to cauterize the other vas deferens. The body wall and skin were sewn up using wound clips. The sterile males were then placed into a clean cage on a warming plate until they recovered from the anesthetic. After 14 days of recovery, a female ICR was mated with the sterile male to verify the male’s inability to fertilize. If the female was plugged but did not become pregnant, it suggested that the vasectomy operation was successful.

#### Mouse Genotyping

The chimeric F0 mice were genotyped by PCR and Sanger sequencing using specific primer pairs. In brief, a small piece of the mouse tail was snipped into an Eppendorf tube. The tail was lysed in 30 μl of Solution A (25 mM NaOH) at 95°C for 20 min and neutralized with 30 μl of Solution B (25 mM Tris-HCl pH 6.8). The 2 μl of the neutralized tail lysis solution was used as a template to screen for the targeted mutations by PCR (95°C, 3 min; 95°C, 15 s, 58°C, 15 s, 72°C, 15 s for 40 cycles; and a final extension at 72°C, 5 min) using specific primers (geno_226228_F and geno_226228_R) (*Table S1*). All of the PCR products were sent for Sanger sequencing to exam the genotype of the chimeric F0 mice. We obtained one chimeric mouse with the precise CTCF ADA mutation and 14 chimeric mice with random indels out of a total of 61 P0 chimeric mice. The targeted chimeric mouse was then crossed with WT mice to obtain heterozygous F1 mice. From a total of 5 litters, we obtained two heterozygous CTCF ADA F1 mice out of a total of only 19 pups. The primer sequences used for mouse genotyping were listed in *Table S1*.

#### RNA-seq

RNA-seq was carried out as previously described with minor modifications (22). The P0 pups or adult mice were decapitated (according to the guidelines approved by IACUC of Shanghai Jiao Tong University). The skin and skull of the head were carefully removed, and the brain was placed in the 35-mm dish filled with ice-cold PBS. The cortical or olfactory bulb tissues were then dissected out using fine forceps and immediately transferred to a new microcentrifuge tube containing 500 μl of ice-cold TRIzol Reagent (Invitrogen 15596018). The sample was homogenized using a pestle and then centrifuged at 12,000 × g for 10 min at 4°C. The supernatant was transferred into a new microcentrifuge tube, and 100 μl of chloroform was added. The sample was then incubated at room temperature for 3 min after vigorous shaking for 15 sec. After centrifuging at 12,000 × g for 15 min at 4°C, the upper aqueous phase was carefully transferred into a new RNase-free microcentrifuge tube, followed by the addition of 250 μl of isopropanol. The mixture was inverted several times and incubated at room temperature for 10 min. After centrifuging at 12,000 × g for 10 min at 4°C, the supernatant was discarded, and the RNA pellet was washed once with 1 ml of freshly-made 75% ethanol, air-dried for 5 min, and dissolved in 30 μl of nuclease-free water. The concentration of RNA was measured using the spectrophotometer (NanoDrop 2000). The high-quality RNA, with an A260/A280 ratio of ∼2.0, is subjected to library preparation.

The polyadenylated mRNA was isolated using oligo(dT) coupled to magnetic beads (Vazyme N401) from 1 μg of total RNA and fragmented by heating at 94°C for 8 min. The first cDNA strand was synthesized by reverse transcription with a random primer (N6). After synthesizing the second strand DNA, the Y-shaped adapter-HTS was ligated to both ends (*Table S1*). The ligated DNA was cleaned by AMPure XP Beads (Beckman A63881). Finally, the RNA-seq library was synthesized through PCR (98°C, 30 s; 98°C, 10 s, 60°C, 30 s, 72°C, 30 s for 10 cycles; and a final extension at 72°C, 5 min) by combining the reagents (2.5 μl of P5 primer (*Table S1*), 2.5 μl of P7 primer (*Table S1*), and 25 μl of VAHTS HiFi Amplification Mix (Vazyme NR604)). All RNA-seq libraries were sequenced on an Illumina NovaSeq 6000 platform.

#### ChIP-seq

Chromatin immunoprecipitation was carried out as previously described with minor modifications (22). Briefly, cortical tissues were dissected out using fine forceps and immediately transferred to a dish containing ice-cold NeuroBasal medium (Gibco 10888022). The tissues were then torn into pieces using fine forceps and transferred to an Eppendorf tube containing 700 μl of 0.25% trypsin-EDTA. The sample was incubated at 37°C for 15 min with continuous shaking at 300 rpm. The trypsin digestion of tissues was quenched by the addition of 700 μl of 10% FBS in the PBS buffer. After filtering the dissociated tissues into a single-cell suspension, the volume was adjusted to 5 ml using 10% FBS in the PBS for crosslinking. The single cells were crosslinked by adding 335 μl of 16% formaldehyde (Thermo Scientific 28906) and incubated at room temperature for 10 min. The reaction was then quenched by the addition of 500 μl of 2 M glycine (Invitrogen 15527013). After centrifuging, the cell pellets were washed twice with 1 ml of ice-cold 1 × PBS (containing 1 × protease inhibitors (Roche 04693116001)). The washed cell pellets were then lysed twice with 1 ml of ice-cold Buffer I (10 mM Tris-HCl pH 7.5, 0.1% sodium deoxycholate (Sigma-Aldrich 30970), 0.15 M NaCl (Sigma-Aldrich S3014), 0.1% SDS (Sigma-Aldrich 71736), 1% Triton X-100 (Sigma-Aldrich T8787), 1 mM EDTA (Sigma-Aldrich E1644), and 1 × protease inhibitors (Roche 04693116001)) at 4°C for 10 min with continuous rotations at 9 rpm. The lysed cells were sonicated with 33% power at a train of 30-sec sonication with 30-sec interval for 30 cycles using the Sonics Vibra-Cell to obtain DNA fragments ranging from 200 to 500 bp. The insoluble debris was discarded by centrifuging at 14,000 × g for 10 min at 4°C. The supernatant cell lysate was initially pre-cleared with 50 μl of protein A/G-agarose beads (Millipore 16-157) for 2 h at 4°C with gentle rotations. The sample was centrifuged at 2,000 × g for 1 min, and the supernatant was then transferred to a new microcentrifuge tube. Finally, an antibody specifically against CTCF (Millipore 07-729) or RAD21 (Abcam ab992) (*Table S2*) was added to the supernatant for immunoprecipitation at 4°C overnight with slow rotations.

On the following day, the antibody-protein-DNA complexes were purified with 50 μl of protein A/G agarose beads by incubating at 4°C for 3 h with slow rotations. The beads were washed once with 1 ml of ice-cold Buffer I, once with 1 ml of ice-cold Buffer I with 0.4 M NaCl, thrice with ice-cold Buffer I without NaCl, and finally with 1 ml of ice-cold LiCl buffer (0.5 M LiCl (Sigma-Aldrich L4408), 50 mM Tris-HCl pH 7.5, 1 mM EDTA, 1% NP-40 (Sigma-Aldrich I8896), 0.7% sodium deoxycholate), by incubating at 4°C for 10 min with slow rotations, respectively. The crosslinked complexes were eluted from the beads using 400 μl of elution buffer (50 mM Tris-HCl pH 8.0, 10 mM EDTA, 1% SDS) at 65°C for 1 h while shaking at 1,000 rpm, followed by reverse crosslinking by incubating at 65°C overnight while shaking at 1,000 rpm. After the addition of 400 μl of phenol-chloroform, the DNA was extracted by vigorous vortexing and centrifuging at 17,000 × g for 15 min at room temperature. After ethanol precipitation, the DNA pellet was resuspended in 50 μl of 10 mM Tris-HCl pH 7.5. The concentration of the DNA was measured using Qubit fluorescent dye (Vazyme EQ121-01). The purified DNA was subjected to high-throughput library preparation following the method described above in the RNA-seq section. All ChIP-seq libraries were sequenced on an Illumina NovaSeq 6000 platform.

#### In situ Hi-C

*In situ* Hi-C experiments were carried out following the previous protocol with minor modifications (22,54). Briefly, after single-cell isolation from the cortical tissues, one million cells were cross-linked with 1% formaldehyde (Thermo Scientific 28906) at room temperature for 10 min with slow rotations. The crosslinked cells were washed twice with 1 × PBS (Gibco 70011044) and lysed twice with 250 μl of ice-cold lysis buffer (10 mM Tris-HCl pH 8.0, 10 mM NaCl, 0.2% NP-40) for 10 min at 4°C to obtain nuclei. Subsequently, the nuclei were permeabilized in 50 μl of 0.05% SDS at 62°C for 10 min, then quenched by the addition of Triton X-100, and digested with 100 U of *Mbo*I (NEB R0147L) in a total volume of 250 μl overnight at 37°C with shaking at 900 rpm. After heat inactivating *Mbo*I at 62°C for 20 min, the digested DNA ends were filled in and labeled by adding a mixture that included DNA Polymerase I, Large (Klenow) Fragment (NEB M0210S), biotin-14-dATP (Thermo 19524016), as well as dCTP, dTTP, and dGTP (NEB N0446S). The reaction was then incubated at 37°C for 1 h.

After labeling the ends of restriction fragments, the digested nuclei were proximally ligated with 2 μl of T4 DNA ligase (NEB M0202S) at room temperature for 4 h. The ligated chromatin was digested using 4 μl of 20 mg/ml proteinase K (Invitrogen AM2546) at 55°C for 4 h and then reverse crosslinked overnight at 68 °C overnight. The DNA was purified through phenol-chloroform extraction followed by ethanol precipitation as described above. After precipitation, the DNA was dissolved in 130 μl of 10 mM Tris-HCl pH 8.0. Sonication of the DNA was carried out using a Bioruptor system at high intensity levels. The process consisted of 15 cycles, each comprising 30-sec sonication followed by 30-sec interval. DNA fragments of 300–500 bp were selected using AMPure XP beads (Beckman A63881) and subsequently captured using streptavidin beads (Thermo 11206D) to enrich the biotin-labeled DNA fragments. The biotin-DNA on the beads was stringently washed twice with 2-min of vortices at 55°C. The purified DNA was used as a template for library preparation as described above. The library was then subjected to sequencing on the MGI-T7 platform via the PE150 sequencing strategy.

#### QHR-4C

Quantitative high-resolution chromosome conformation capture copy (QHR-4C) experiments were performed as previously described (6) with minor modifications. In brief, five million cells were harvested as described above. The cells were resuspended in 10 ml of MEM medium, and then crosslinked with 2% formaldehyde (Thermo Scientific 28906) at room temperature with slow rotations for 10 min. The reaction was quenched by the addition of 700 μl of 2 M glycine (Invitrogen 15527013). The fixed cells were spun down at 900 × g for 5 min at 4°C and then washed twice with 1 ml of ice-cold PBS (Gibco 70011044). Cells were lysed twice with 1 ml of ice-cold 4C permeabilization buffer (50 mM Tris-HCl pH 7.5, 150 mM NaCl, 5 mM EDTA, 0.2% SDS, 0.5% NP40, 1% Triton X-100, and 1 × protease inhibitors) with slow rotations for 10 min at 4°C. Cells were then resuspended in 73 μl of nuclease-free water, followed by the addition of 10 μl of 10 × *Dpn*II buffer (NEB R0543L) and 2.5 μl of 10% SDS, and incubation at 37°C with constant shaking at 900 rpm. After 1 h, 12.5 μl of 20% Triton X-100 (Sigma-Aldrich T8787) was added to the sample, and the sample was incubated at 37°C for an additional hour with constant shaking at 900 rpm. Next, 8 μl of *Dpn*II (NEB R0543L) was added to the sample, followed by an overnight incubation at 37°C with constant shaking at 900 rpm. The enzyme was heat-inactivated at 65°C for 20 min. The sample was centrifuged at 2,000 × g for 10 min, and the supernatant was carefully removed to not disturb the cell pellets. The cell pellets were then resuspended with 1,600 μl of nuclease-free water. The digested nuclei were ligated with 2 μl of T4 DNA ligase (NEB M0202S) in a total ligation volume of 2 ml at 16°C for 12 h.

The next day, the ligated chromatin was digested with 8 μl of 10 mg/ml Protease K (Invitrogen AM2546) at 65°C for 2 h. The DNA was extracted using phenol-chloroform extraction, followed by ethanol precipitation as described above. The DNA pellet was then dissolved in 50 μl of nuclease-free water and sonicated using a Bioruptor system with a low energy setting. Sonication was performed in cycles of 30-sec on and 30-sec off for 12 cycles to obtain DNA fragments ranging from 200-600 bp. The concentration of the sonicated DNA was measured using the Qubit fluorescent dye (Vazyme EQ121-01). Next, 5 μg of DNA was used as a template for linear amplification with a specific 5’ biotin-tagged primer complementary to the viewpoint. The linear amplification was conducted in a total volume of 100 μl, including 50 μl of 2 × Buffer, 0.8 μl of dNTP, 1.6 μl of Phanta (Vazyme P505-d1), and 1 μl of 1 μM 5’ biotin-tagged primer (*Table S1*), using the program (95°C, 2 min; 95°C, 15 s, 58°C, 25 s, 72°C, 1 min 30 s for 80 cycles; and a final extension at 72°C, 5 min). The amplified products were denatured at 95°C for 5 min and then chilled on ice to obtain biotinylated single-stranded DNA (ssDNA). The biotinylated ssDNA was ligated with 2 μl of 50 μM Adapter-4C (*Table S1*) adapters in a total volume of 45 μl for 12 h at 16°C and purified by streptavidin magnetic C1 beads (Invitrogen 65001). The QHR-4C libraries were generated via PCR amplification (94°C, 2 min; 94°C, 10 s, 60°C, 15 s, 72°C, 1 min for 19 cycles; and a final extension at 72°C, 5 min) using Phanta high-fidelity DNA polymerase (Vazyme P505-d1). The amplified PCR products were purified using the High-Pure PCR Product Purification kit (Roche 11732676001) and sequenced on an Illumina NovaSeq 6000 platform. The primers for QHR-4C are listed in *Table S1*.

### Bioinformatics analyses

#### RNA-seq analysis

Raw FASTQ data files were mapped to the mouse reference genome (GRCm38/mm10) using the HISAT2 (v2.2.1) (55) with default parameters. Subsequently, Cufflinks (v2.2.1) (56) was employed to estimate transcript expression levels in FPKM (Fragments Per Kilobase of exon per Million mapped fragments). Gene expression analyses were conducted with a minimum of three replicates, presenting data as mean ± S.D. Statistical significance of gene expression changes was determined using unpaired Student’s *t*-test in GraphPad Prism (v8.4.2) and R (v4.1.1), with significance levels denoted as follows: ‘*’ for *P* ≤ 0.05, ‘**’ for *P* ≤ 0.01, and ‘***’ for *P* ≤ 0.001.

For the genome-wide analyses of differentially expressed genes (DEGs), featureCounts (v2.0.1) (57) was used to generate the raw count matrix. The count matrix was then assayed using the R package DESeq2 (58) to detect DEGs, employing parameters of absolute log_2_FC > 0.5 and *padj* < 0.05. Subsequently, a volcano plot depicting DEGs was generated using the R package ggplot2 (v3.5.0) (59).

Manhattan plots illustrating the localized enrichments of downregulated genes were produced using the R package CMplot (v4.5.1) (27,60). In brief, the mouse reference genome (GRCm38/mm10) was partitioned into non-overlapping 1-Mb bins. Subsequently, the genomic locations of the downregulated genes were intersected with these 1-Mb bins to determine the frequency of hits in each bin. The occurrence frequency of each signal within 1-Mb bins was then modeled using a Poisson distribution. The maximum-likelihood estimator for the λ was determined based on the mean number of occurrences. Subsequently, the Poisson models were employed to compute the probability of observing each signal within every 1-Mb bin. The probabilities were documented within each bin and subjected to CMplot (v4.5.1) (60) to generate Manhattan plots.

Scatterplots showing fold changes of expression levels of *cPcdh* genes as a function of the linear genomic distance between promoters and enhancers were generated using ggplot2 (v3.5.0) (8,59). Briefly, the fold change for each *cPcdh* gene was calculated based on the RNA-seq data. The enhancer element associated with the *Pcdh α*, *β*, and *γ* clusters are *HS5-1*, *HS18-22*, and *HS5-1bL*, respectively. The 5’ untranslated regions (UTRs) of each gene were served as anchor points for calculating the distances between the genes and enhancers. Subsequently, the fold changes for each gene were paired with their respective distances from the enhancers based on gene names. These data were used to create the scatterplots by utilizing ggplot2 (v3.5.0) (59). A smooth curve is added to the scatterplot to facilitate the observation of trends in the data using ggplot2 (v3.5.0) (59).

Bar plots displaying the proportions of mutated bases in high-throughput RNA-seq reads were generated using ggplot2 (v3.5.0) (59). The counts of mutated bases were visualized and summarized using the Integrative Genomics Viewer (IGV) (61). The ratio was computed manually and then plotted using ggplot2 (v3.5.0) (59) to create the bar graphs.

#### ChIP-seq analysis

The raw sequencing data were mapped to the mouse reference genome (GRCm38/mm10) using Bowtie2 (v2.2.5) (62) with default settings. Samtools (v1.15.1) (63) was utilized to convert the Sequence Alignment/Map (SAM) format into a Binary Alignment/Map (BAM) format and to index the resulting BAM files. Subsequently, the BAM files were normalized to RPKM (Reads Per Kilobase per Million mapped reads) for comparative analysis of occupancy levels across samples, utilizing the bamCoverage module of the deepTools suite (64). The bedGraph files, generated from the deepTools program (64), were then uploaded to the UCSC Genome Browser (65) for visualization of genomic regions of interest. Peak calling was conducted using MACS2 (v2.2.7.1) (66) with default parameters.

Boxplots illustrating the ChIP-seq profiles of RAD21 in the *cPcdh* locus were generated using ggplot2 (v3.5.0) (59). Statistical significance of changes of RAD21 enrichment was computed using the R package rstatix (v0.7.0), with significance levels denoted as following: ‘*’ for *P* ≤ 0.05, ‘**’ for *P* ≤ 0.01, and ‘***’ for *P* ≤ 0.001.

Scatterplots showing fold changes of RAD21 enrichments at the *cPcdh* genes as a function of the linear genomic distance between promoters and enhancers were created using ggplot2 (v3.5.0) (59). Briefly, RAD21 enrichments for each *cPcdh* gene were calculated based on the ChIP-seq data by calculating the FPKM of all peaks. The fold changes in RAD21 enrichments in heterozygous mice were then computed by comparing with the WT littermates. Distances between genes and enhancers were calculated using the 5’ untranslated regions (UTRs) of each gene as anchor points. The fold changes in Rad21 enrichments for each gene were paired with their corresponding distances from the enhancers based on gene names.

#### Hi-C analysis

Raw FASTQ reads were mapped to the mouse reference genome assembly (GRCm38/mm10) using the HiC-Pro pipeline (67) (*Table S3*). The resulting allValidPairs files were converted to the hic format files using the hicpro2juicebox script from the HiC-Pro toolkit, enabling analysis with Juicer tools (68). Subsequently, the hic format was converted to cooler format using hic2cool (v0.8.3). To ensure accuracy, the contact matrix was normalized to a contact depth of 100 million based on *cis*-interactions using the Knight-Ruiz (KR) method.

The Hi-C contact maps for targeted genomic regions were generated using the fancplot module in FAN-C toolkits (69), and the differential contact heatmaps were constructed using Seaborn, a Python library. At a 100-Kb resolution, the Hi-C *cis*-Eigenvector 1 values and Pearson’s correlation matrix were calculated utilizing the Juicer software (68). Subsequently, the resulting *cis*-Eigenvector 1 values were converted into the bedGraph format for visualization in the UCSC genome browser (65). Scatterplots with a heat density were depicted using the R package ggpointdensity (v0.1.0).

TADs were called using the Hidden Markov Models (HMM) approach, as reported in a previous study (70). The dense matrix was subjected to DI_from_matrix script to calculate the Directionality Index (DI) score. The DI score was then utilized to predict the hidden states of all bins via HMM models. Subsequently, bins with consecutive identical states were merged into regions. Regions containing less than 3 bins or with a median posterior bin probability below 0.99 were excluded from the analysis using a Perl script. Ultimately, TADs were defined as regions extending downstream from an upstream boundary and upstream from a downstream boundary. The DI score was transformed into the bedGraph format to visualize in the UCSC genome browser.

To identify loops in the contact matrix, Chromosight (v1.6.3) (71) was employed to detect chromatin loops with a genomic distance greater than 4 Kb. Subsequently, specific loops were analyzed using the Aggregate Peak Analysis (APA) with cooltools (v0.6.1) (72) and coolpup.py (v1.1.0) (73).

#### QHR-4C analysis

Raw FASTQ files were mapped to the mouse reference genome (GRCm38/mm10) using Bowtie2 (v2.2.5) (62) with default parameters. The reads per million (RPM) values were calculated using r3Cseq (74) program (v1.20). Bedgraph files generated by r3Cseq (v1.20) were visualized in the UCSC genome browser.

Scatterplots depicting fold changes of contact probabilities of *cPcdh* genes as a function of the linear genomic distance between target variable promoters and their respective enhancers were created using ggplot2 (v3.5.0). Interaction values were assigned to the variable exons of each member of the *cPcdh* genes, and the fold changes in contact probabilities upon CTCF ADA mutation were quantified. Gene-enhancer distances were computed utilizing the 5’ untranslated regions (UTRs) of each gene as reference points. The fold change data for each gene were matched with distances from the enhancers based on gene names. These datasets were employed in ggplot2 (v3.5.0) to produce scatterplots, with a smoothing curve included to facilitate the visualization of data trends.

## Data availability

High-throughput sequencing files have been deposited in NCBI’s Gene Expression Omnibus (GEO) database under the accession numbers GSE285577, GSE285578, GSE285579, and GSE287859 respectively. Hi-C raw data have been deposited in SRA database under the accession number PRJNA1213240. Raw WB data have been deposited to Zenodo: DOI: 10.5281/zenodo.14672244.

## Acknowledgments

We would like to express our gratitude to all members of our laboratory for their insightful discussions. This work was supported by grants to Q.W. from the National Natural Science Foundation of China (32330016) and the National Key R&D Program of China (2022YFC3400200).

## CRediT authorship contribution statement

Q.W. conceived and supervised the project. Y.Z. performed experiments and analyzed data. Y.Z. and Q.W. wrote the manuscript.

---

**Figure S1.**
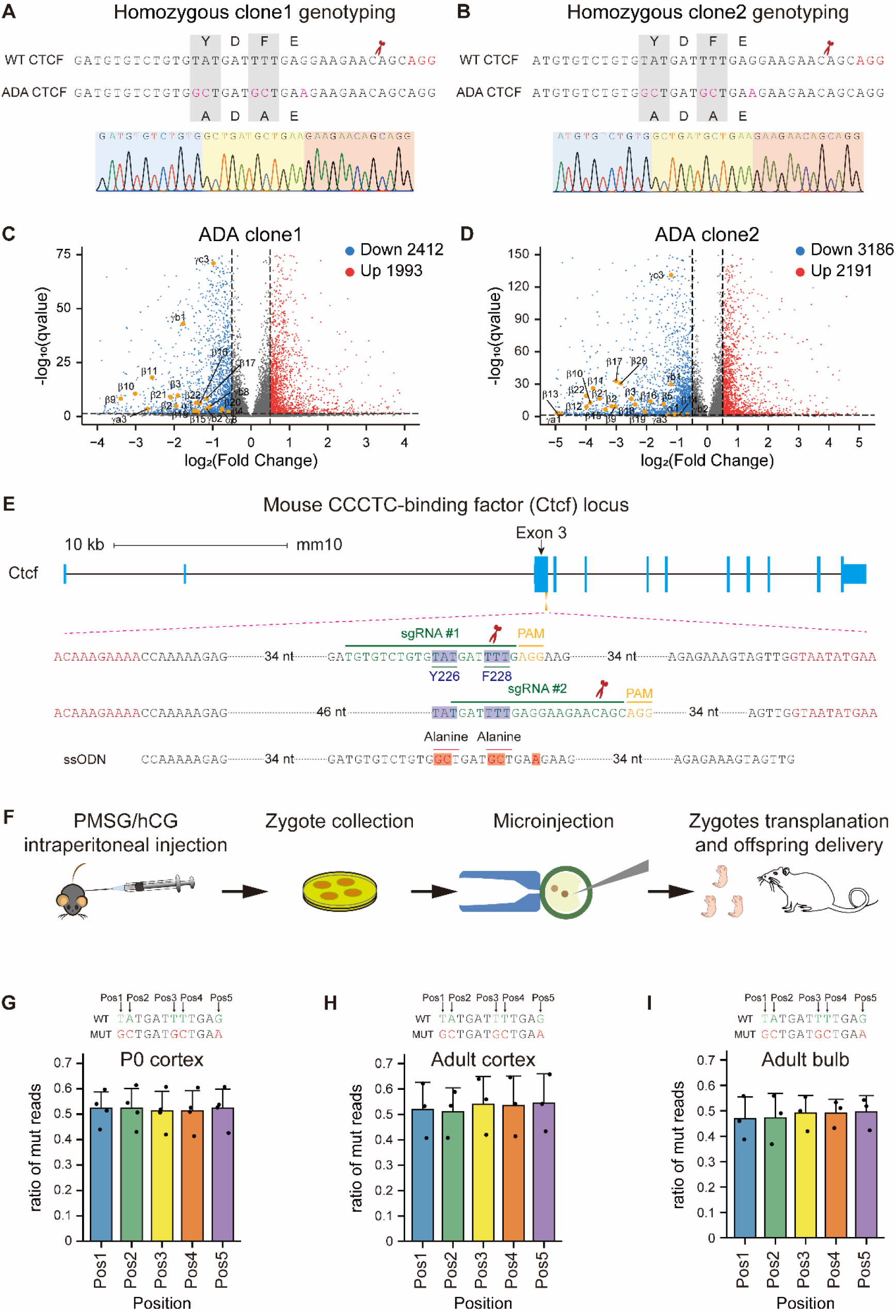
Generation of models of CTCF ADA mutation. *A* and *B,* Genotyping of two independent homozygous single-cell clones of CTCF ADA mutation. *C* and *D,* Volcano plots of differentially expressed genes in each of the two independent *N2a* cell clones. The downregulated *cPcdh* genes are marked with yellow dots. *E,* Schematic of CRISPR-mediated DNA editing with a single stranded oligodeoxynucleotide (ssODN) donor for CTCF ADA mutation in mice. *F,* Outline of the procedure for generating CRISPR-mediated site-directed mutagenesis in the N-terminal domain of CTCF in mice. *G*-*I,* Bar plots showing ratios of mutated bases in high-throughput RNA-seq reads from the heterozygous P0 (n=4) (*G*) and adult (n=3) (*H*) cortical, as well as olfactory bulb (n=3) (*I*) tissues. nt, nucleotide. sgRNA, single guide RNA. PAM, protospacer adjacent motif. ssODN, single-stranded oligodeoxynucleotide. PMSG, pregnant mare serum gonadotropin. hCG, human chorionic gonadotropin. Pos, position. Data are mean ± S.D from at least three biological replicates.

**Figure S2.**
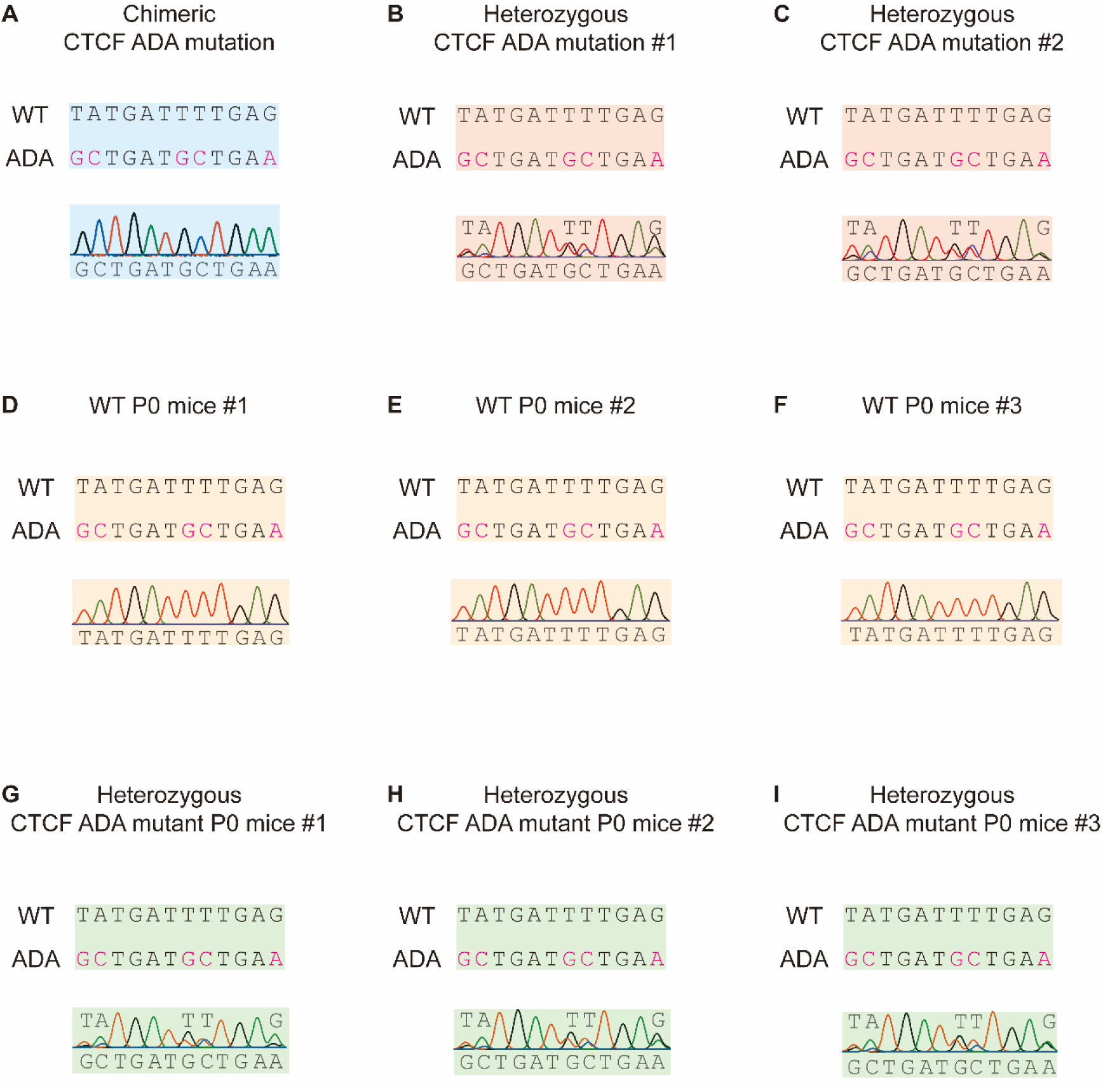
Genotyping of mouse models of CTCF ADA mutation. *A,* Genotyping of the chimeric CTCF ADA mouse by TA cloning and Sanger sequencing. *B* and *C,* Genotyping of the heterozygous CTCF ADA mice by Sanger sequencing. *D*-*I,* Genotyping of a litter of three P0 wild-type mice and three P0 heterozygous CTCF ADA mice by Sanger sequencing.

**Figure S3.**
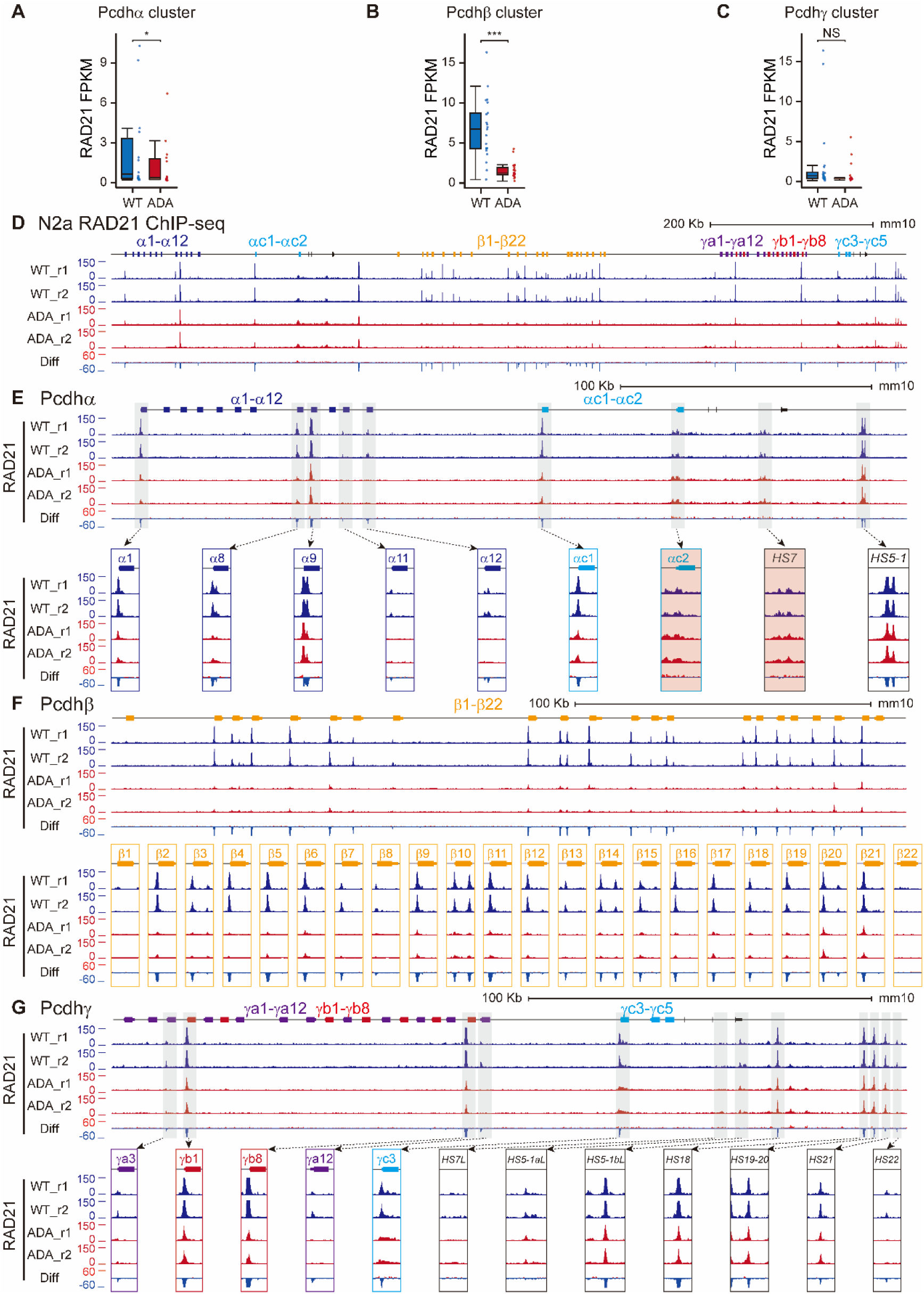
Decreased enrichments of RAD21 at the promoter and enhancer regions of the *cPcdh* genes in *N2a* cells. *A*-*C,* Quantification of alterations of RAD21 enrichments at the *Pcdh α* (*A*), *β* (*B*), and *γ* (*C*) clusters in the cellular model. Statistical significance of changes of RAD21 enrichment was computed using the R package rstatix (v0.7.0), with significance levels denoted as following: ‘*’ for *P* ≤ 0.05, ‘**’ for *P* ≤ 0.01, and ‘***’ for *P* ≤ 0.001, using paired two-tailed Student’s *t*-test. *D,* RAD21 ChIP-seq profiles at the *cPcdh* locus in CTCF ADA *N2a* clones compared to the WT clone. *E*-*G,* Close-up of RAD21 profiles at the *Pcdh α* (*E*), *β* (*F*), and *γ* (*G*) clusters showing decreased enrichments of RAD21 at *cPcdh* variable exons and enhancers. Note that as internal controls with no CTCF site, there do not appear decreased RAD21 enrichments at *αc2* and *HS7*. r1, replicate 1. r2, replicate 2.

**Figure S4.**
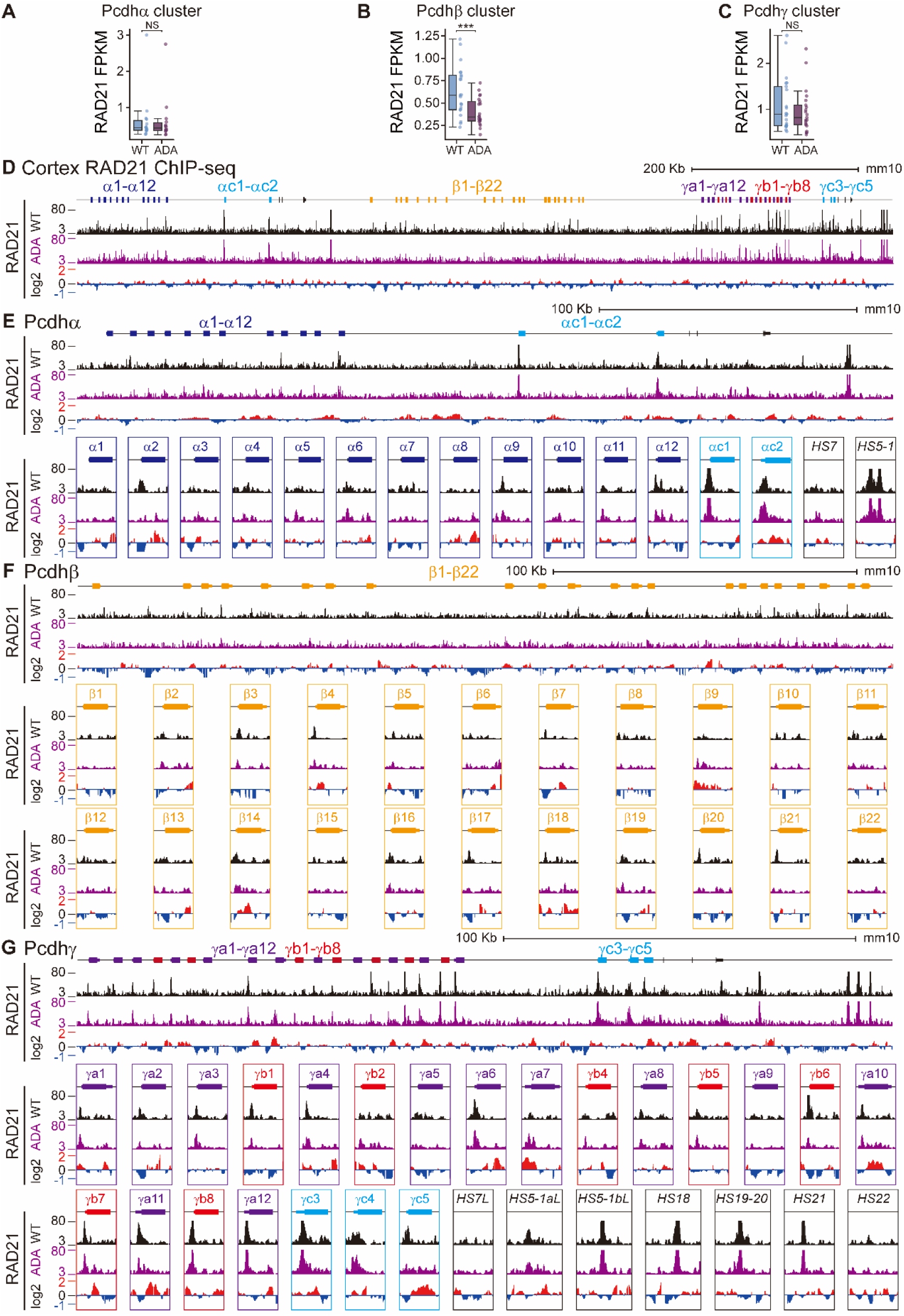
Decreased enrichments of RAD21 at the promoter and enhancer regions of the *cPcdh* locus in the mouse brain. *A*-*C,* Quantification of alterations of RAD21 enrichments at the *Pcdh α* (*A*), *β* (*B*), and *γ* (*C*) clusters in the mouse model. Statistical significance of alterations of RAD21 enrichment was calculated using paired two-tailed Student’s *t*-test, with significance levels denoted as following: ‘*’ for *P* ≤ 0.05, ‘**’ for *P* ≤ 0.01, and ‘***’ for *P* ≤ 0.001. *D,* RAD21 ChIP-seq profiles at the *cPcdh* locus in heterozygous CTCF ADA mice compared to WT littermates. *E*-*G,* Close-up of RAD21 profiles at the *Pcdh α* (*E*), *β* (*F*), and *γ* (*G*) clusters showing decreased enrichments of RAD21 at *cPcdh* variable exons and enhancers.

**Figure S5.**
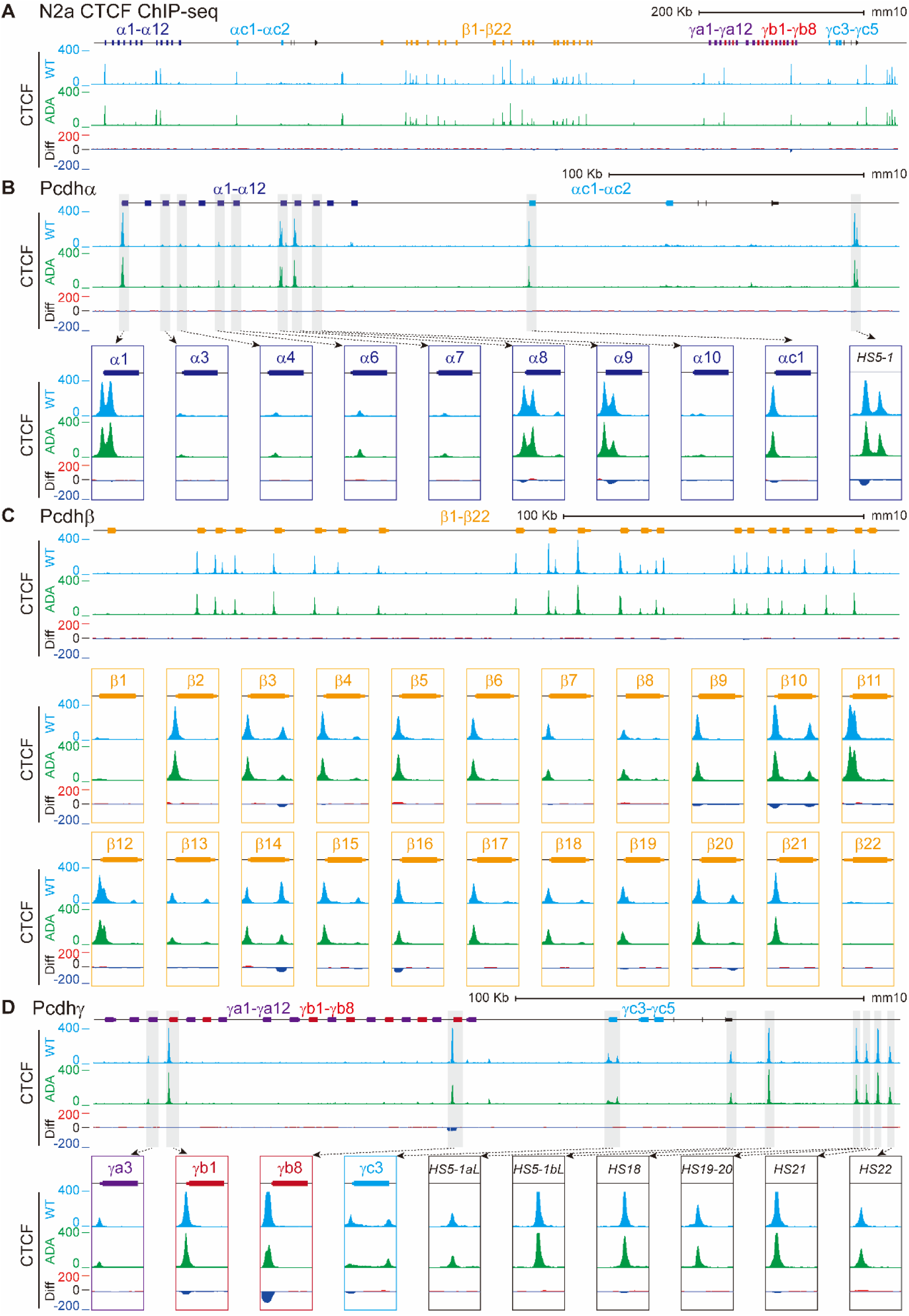
Enrichments of CTCF at the promoter and enhancer regions of the *cPcdh* genes in *N2a* cells. *A,* CTCF ChIP-seq profiles at the *cPcdh* locus in CTCF ADA *N2a* clones compared to the WT clone. *B*-*D,* Close-up of CTCF profiles at the *Pcdh α* (*B*), *β* (*C*), and *γ* (*D*) clusters showing no decrease of CTCF enrichments at *cPcdh* variable exons and enhancers.

**Figure S6.**
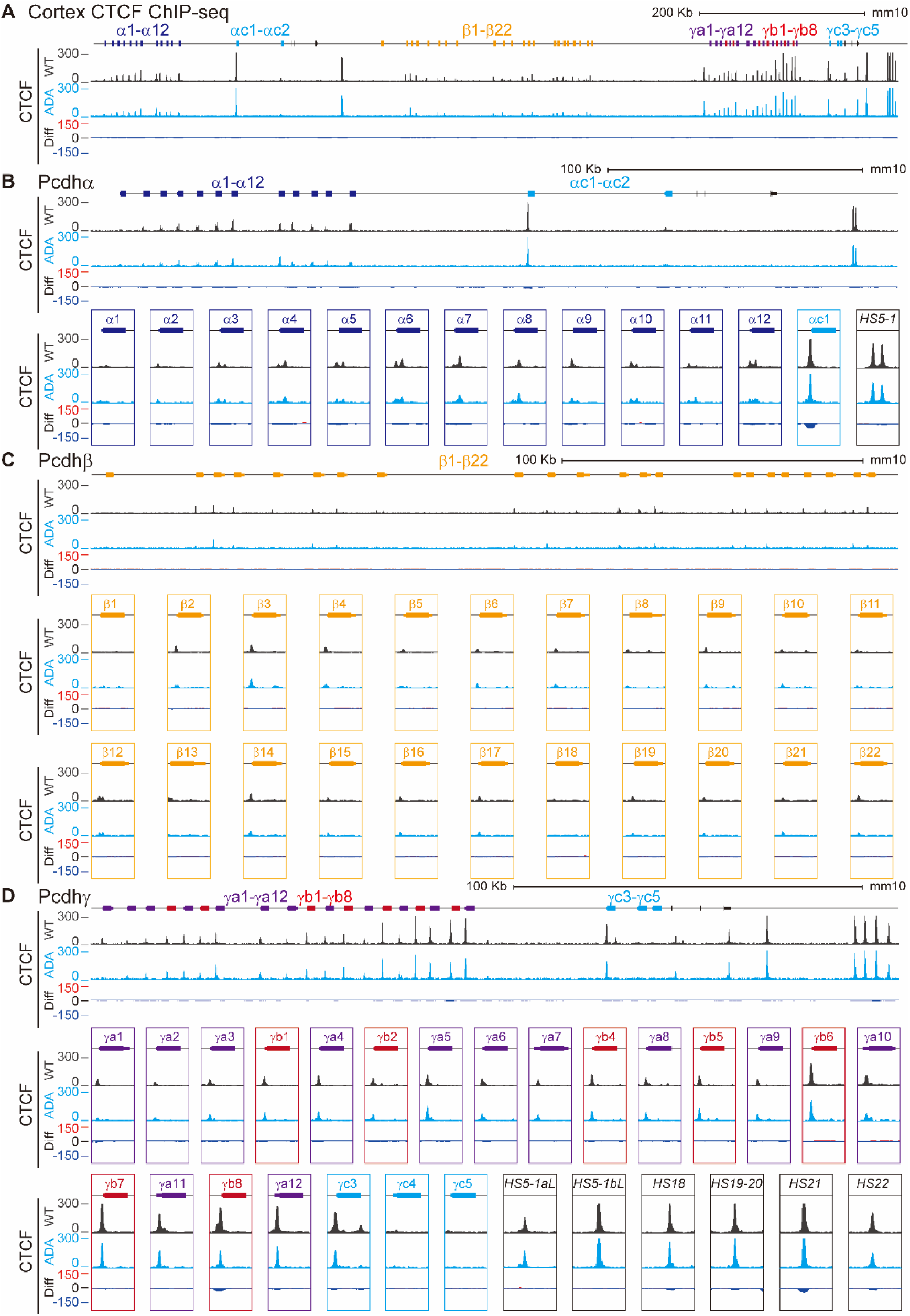
Enrichments of CTCF at the promoter and enhancer regions of the c*Pcdh* locus in the mouse brain. *A,* ChIP-seq profiles of CTCF at the *cPcdh* locus in WT and heterozygous CTCF ADA mice. *B*-*D,* Close-up of CTCF profiles at the *Pcdh α* (*B*), *β* (*C*), and *γ* (*D*) clusters.

**Figure S7.**
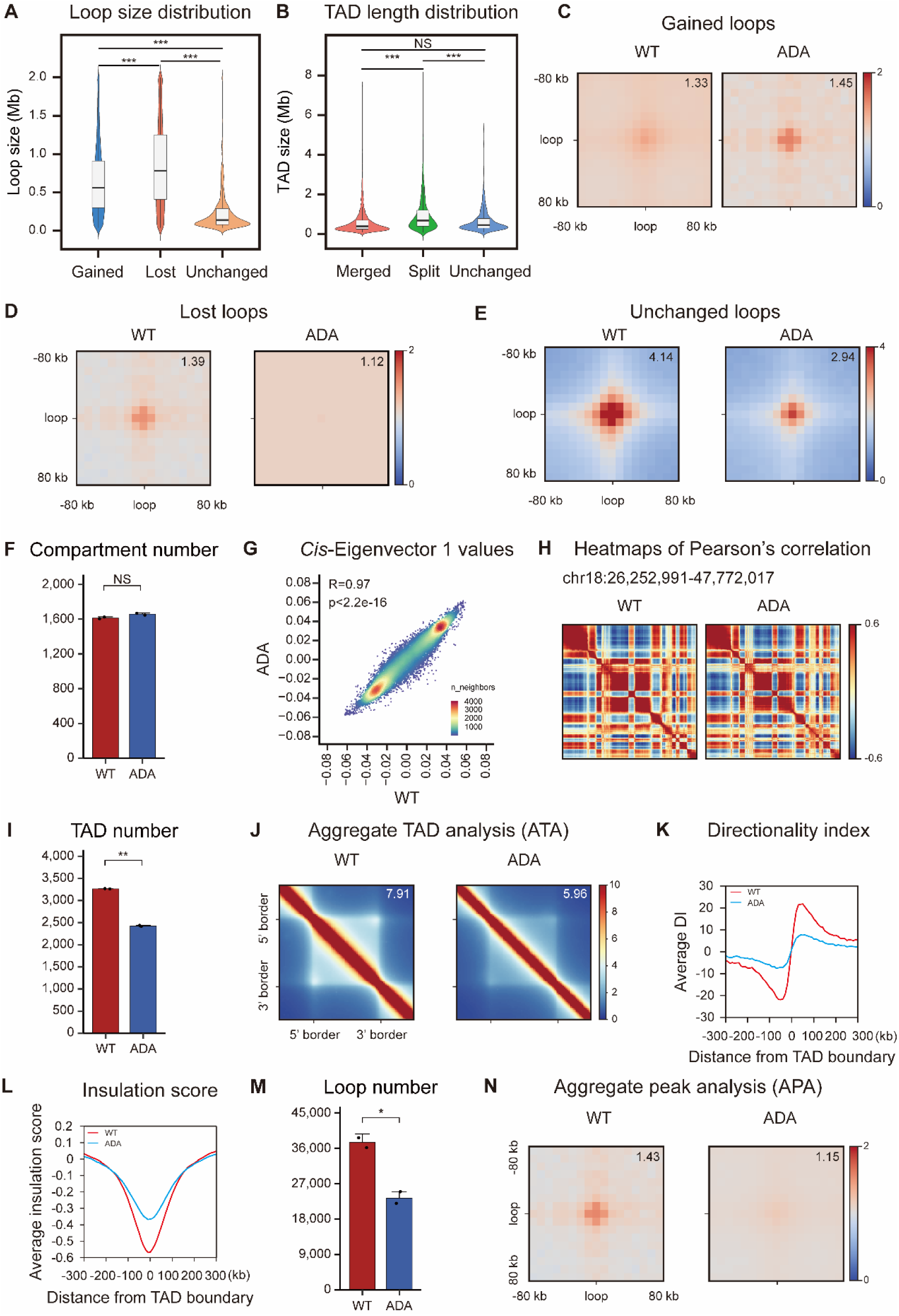
CTCF YDF motif enables cohesin to have a long residence time on chromatin. *A,* Violin plot showing the distribution of loop sizes in gained, lost, and unchanged regions, showing that unchanged loops display the smallest size, whereas lost loops typically tend to have the largest size, and gained loops are in an intermediate range. *B,* Violin plot depicting the distribution of lengths of merged, split, and unchanged TADs, showing that the length of split TADs tends to be significant longer than that of the merged or unchanged TADs. Statistical significance of alterations was calculated using unpaired two-tailed Student’s *t*-test, with significance levels denoted as following: ‘*’ for *P* ≤ 0.05, ‘**’ for *P* ≤ 0.01, and ‘***’ for *P* ≤ 0.001. *C*-*E,* Aggregate peak analysis (APA) of signals of the gained (*C*), lost (*D*), or unchanged (*E*) loops. The APA analysis revealed a significant decrease in loop strength of lost loops, alongside a notable increase in looping strength based on the gained loops upon CTCF ADA mutation. The unchanged loops also exhibited a significant decrease in looping strength, indicating that CTCF ADA mutation destabilizes the extruding cohesin complex on chromatin. In each panel, the APA value is indicated in the upper-right corner. *F,* Bar plot showing numbers of compartment in wild-type and heterozygous CTCF ADA mice (n=2). Data are mean ± S.D from two biological replicates, \**P* < 0.05, \*\**P* < 0.01, \*\*\**P* < 0.001; unpaired two-tailed Student’s *t*-test. *G,* Scatterplot displaying the Pearson’s correlation of *cis*-eigenvector 1 values between wild-type and heterozygous CTCF ADA mice. *H,* Heatmaps of Pearson’s correlation at chr18: 26,252,991-47,772,017 in wild-type and heterozygous CTCF ADA mice, showing an unchanged compartmentalization upon CTCF ADA mutation. *I,* Numbers of TADs in wild-type and heterozygous CTCF ADA mice (n=2). Data are mean ± S.D from two biological replicates, \**P* < 0.05, \*\**P* < 0.01, \*\*\**P* < 0.001; unpaired two-tailed Student’s *t*-test. *J,* Aggregate TAD analysis (ATA) in wild-type and heterozygous CTCF ADA mice, showing a significant decrease of intra-TAD contacts. *K,* Pileup of the directionality index scores in 300-Kb regions centered on all TAD boundaries in wild-type and heterozygous CTCF ADA mice, showing the weakened insulation at TAD boundaries. DI, directionality index. *L,* Averaged insulation scores in 300-Kb regions centered on all TAD boundaries in wild-type and heterozygous CTCF ADA mice. *M,* Hi-C loop number in wild-type and heterozygous CTCF ADA mice (n=2). Data are mean ± S.D from two biological replicates, \**P* < 0.05, \*\**P* < 0.01, \*\*\**P* < 0.001; unpaired two-tailed Student’s *t*-test. *N,* Aggregate peak analysis (APA) depicting the superimposed signal for genome-wide loops estimated from Hi-C data of wild-type and heterozygous CTCF ADA mice, showing global weakening of chromatin loops upon CTCF ADA mutation. In each panel, the APA value is indicated in the upper-right corner.

**Figure S8.**
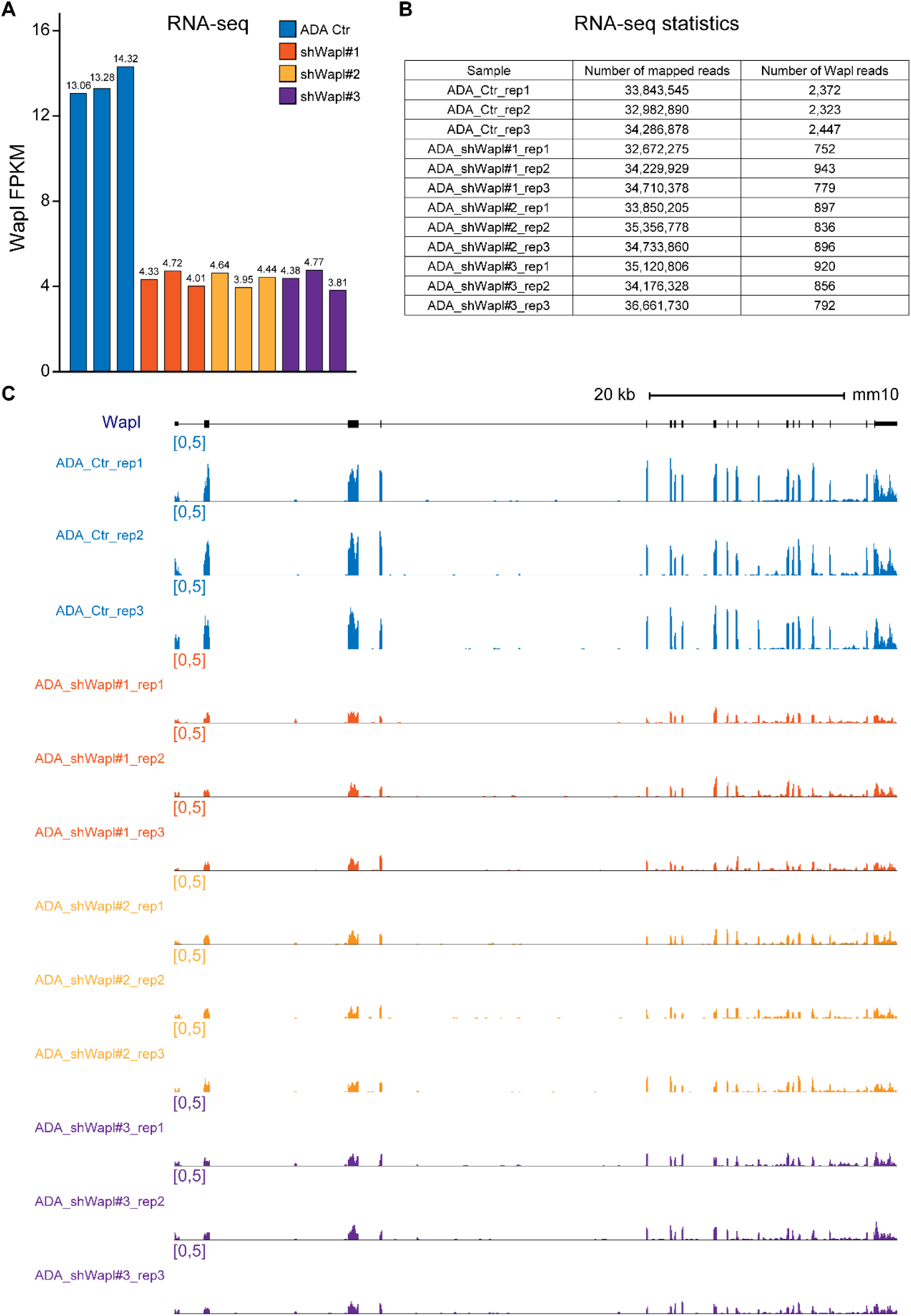
Raw data of Wapl knockdown.

**Table S1.**
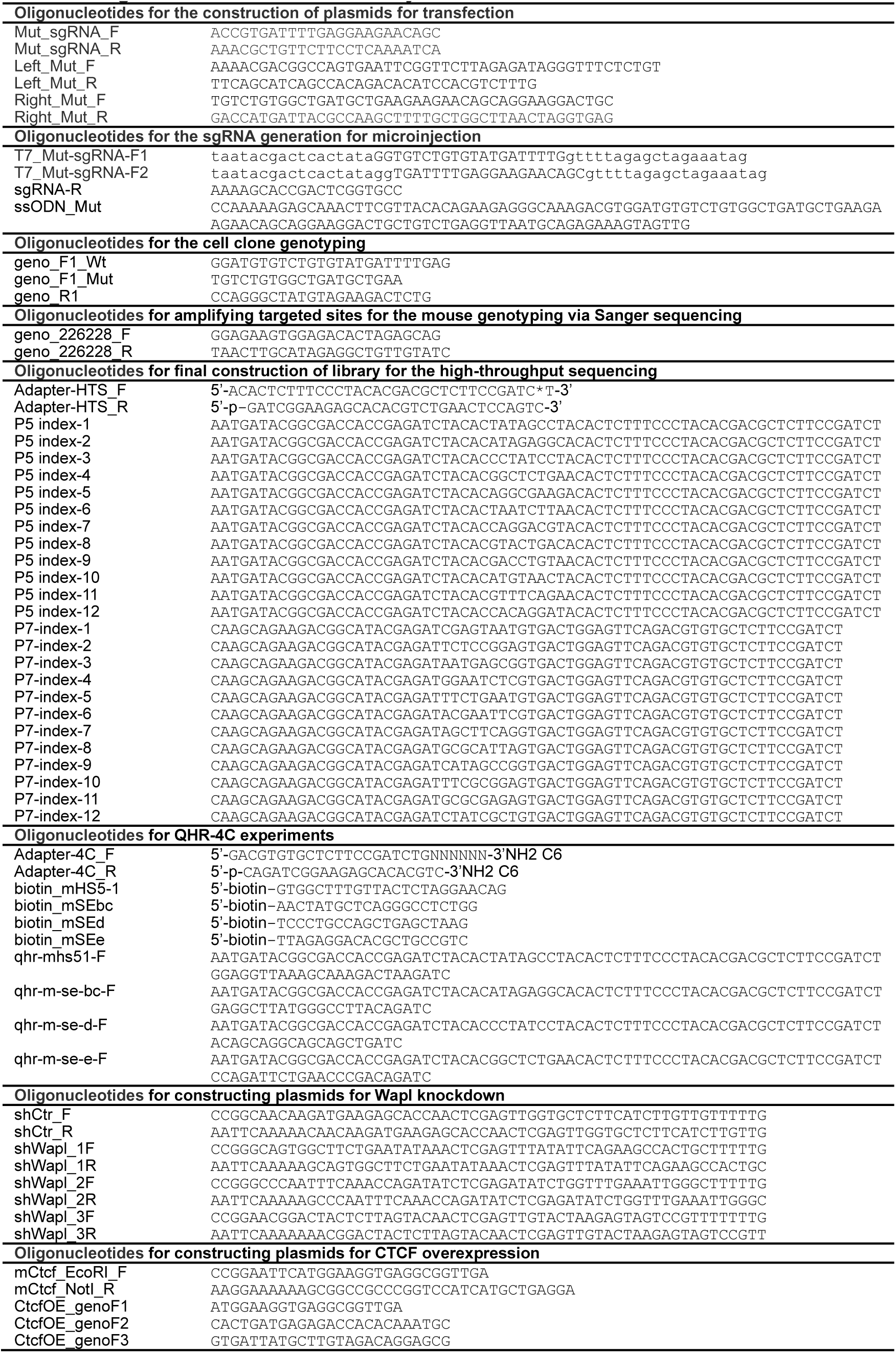
Oligonucleotides used in this study.

**Table S2.**
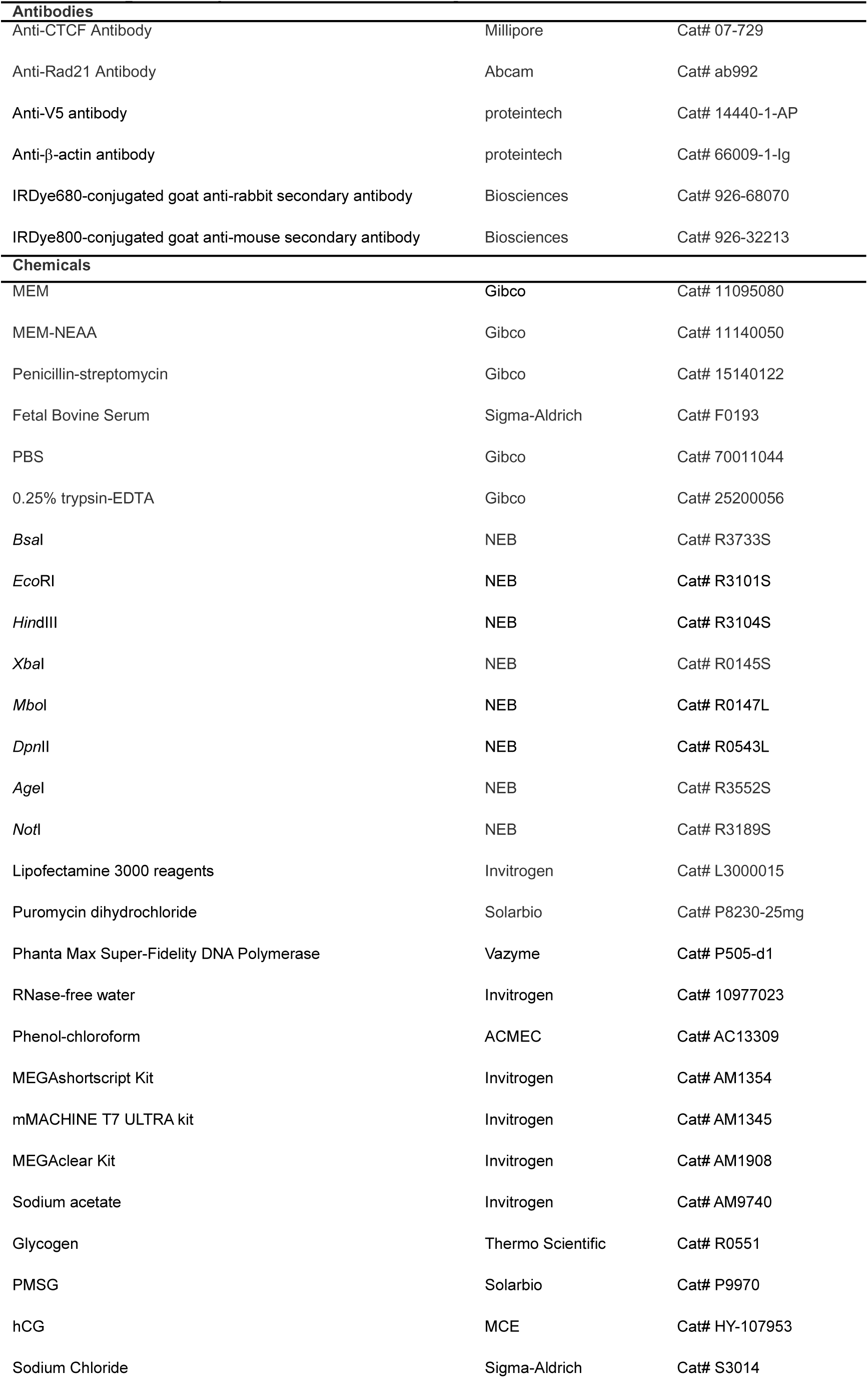
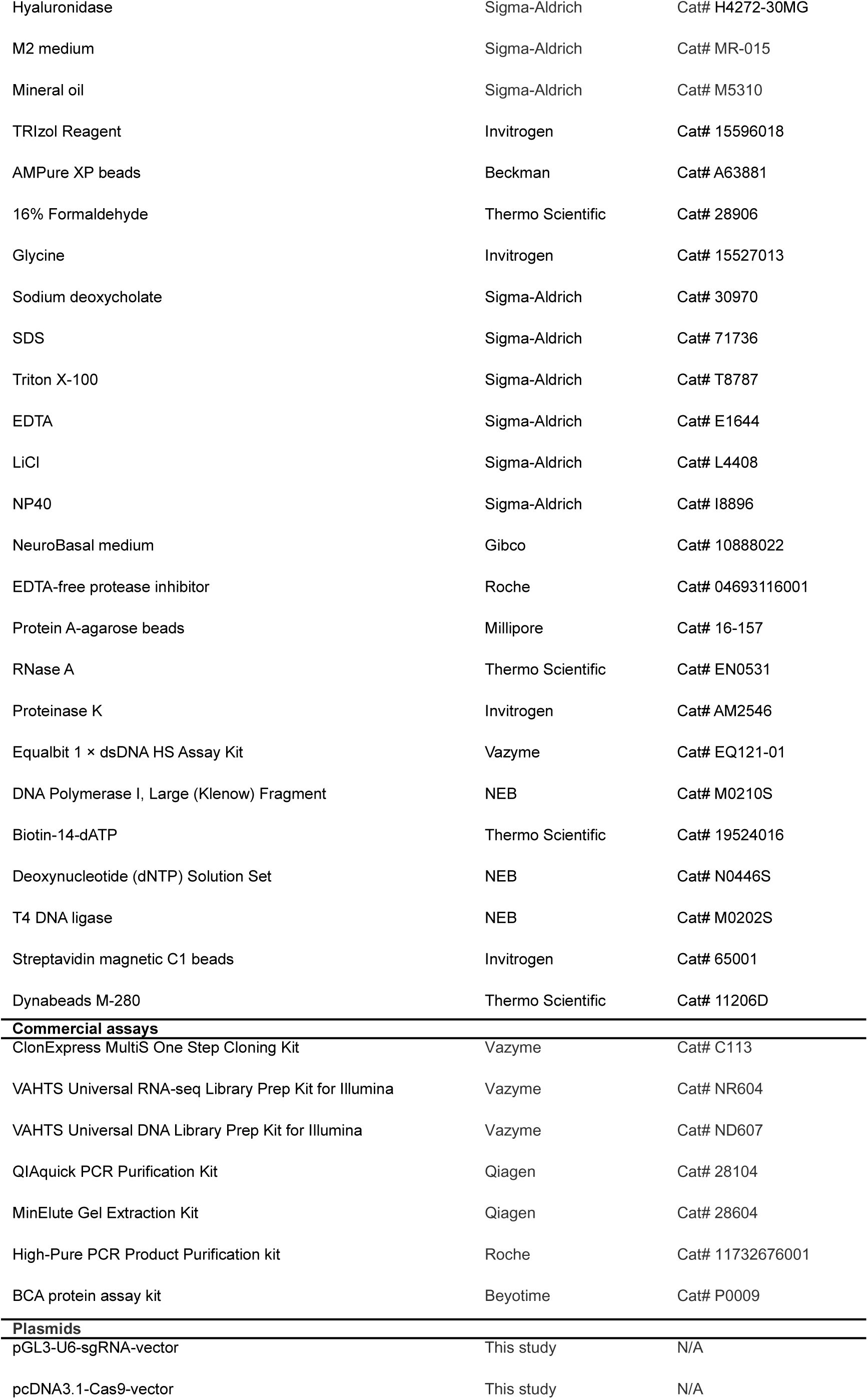
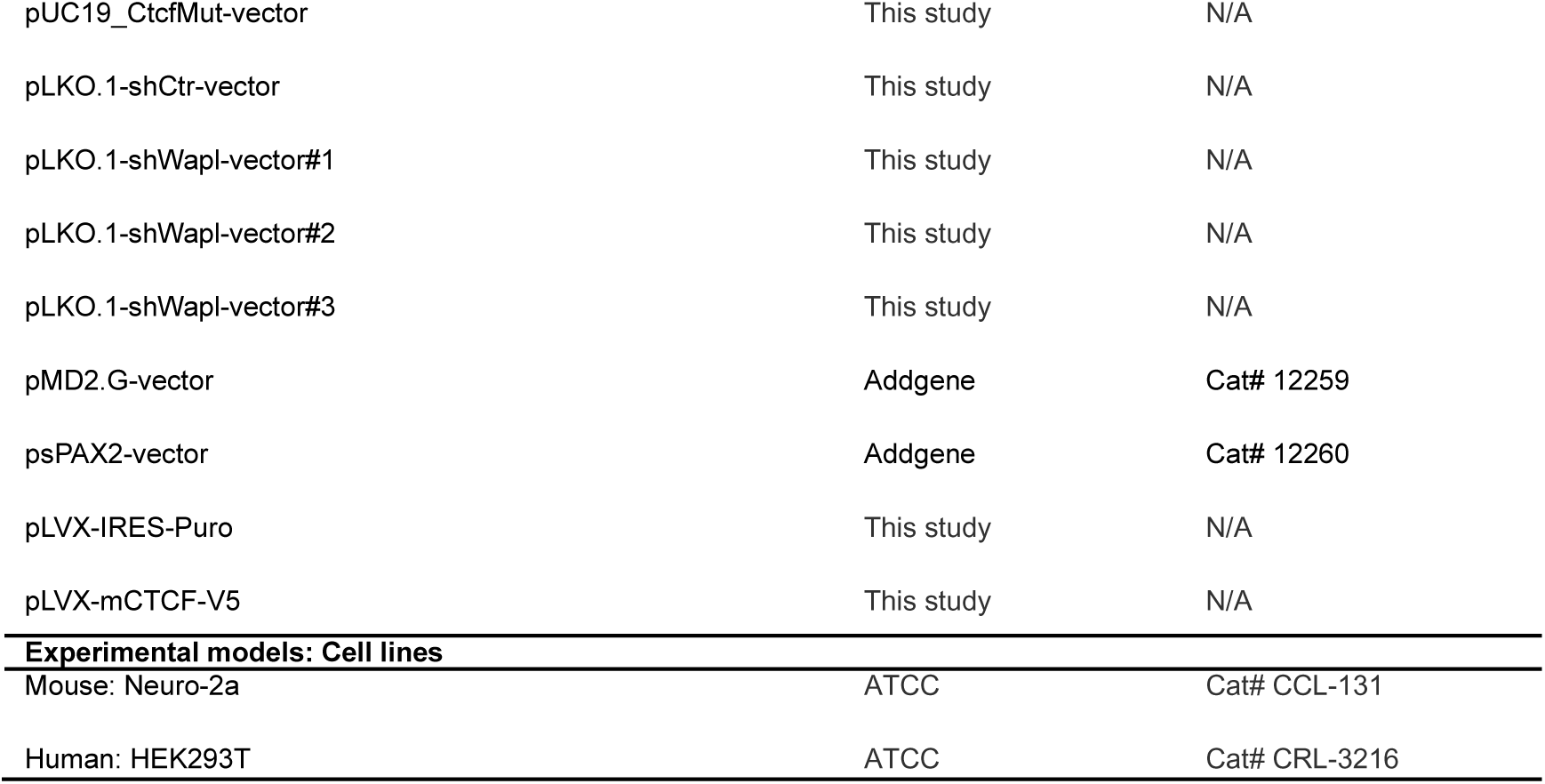
Reagents and plasmids used in this study.

**Table S3.** *In situ* Hi-C statistics.

|  | WT_rep1 | WT_rep2 | ADA_rep1 | ADA_rep2 |
| --- | --- | --- | --- | --- |
| Valid_interaction_pairs | 1,169,250,659 | 1,282,103,567 | 1,056,315,746 | 1,164,695,035 |
| Valid_interaction_pairs_FF | 289,994,816 | 318,174,399 | 263,315,179 | 289,472,201 |
| Valid_interaction_pairs_RR | 289,921,136 | 318,100,330 | 263,312,165 | 289,512,479 |
| Valid_interaction_pairs_RF | 288,683,416 | 316,540,127 | 262,692,179 | 288,544,162 |
| Valid_interaction_pairs_FR | 300,651,291 | 329,288,711 | 266,996,223 | 297,166,193 |
| Dangling_end_pairs | 66,522,829 | 75,925,553 | 46,541,563 | 76,450,325 |
| Religation_pairs | 33,099,080 | 36,340,974 | 16,334,596 | 27,451,754 |
| Self_Cycle_pairs | 3,555,453 | 3,745,368 | 2,968,630 | 2,581,732 |
| Single-end_pairs | 0 | 0 | 0 | 0 |
| Filtered_pairs | 0 | 0 | 0 | 0 |
| Dumped_pairs | 467,118 | 397,900 | 413,642 | 261,100 |
| Valid_interaction | 1,169,250,659 | 1,282,103,567 | 1,056,315,746 | 1,164,695,035 |
| Valid_interaction_rmdup | 1,075,932,537 | 1,176,201,570 | 869,554,809 | 1,019,127,664 |
| Trans_interaction | 373,187,735 | 395,027,651 | 402,791,776 | 490,664,581 |
| Cis_interaction | 702,744,802 | 781,173,919 | 466,763,033 | 528,463,083 |
| Cis_shortRange | 153,890,242 | 171,454,035 | 82,565,133 | 99,359,414 |
| Cis_longRange | 548,854,560 | 609,719,884 | 384,197,900 | 429,103,669 |

